# Shaping dendritic NMDA spikes by timed synaptic inhibition: Implications for the I/O properties of cortical neurons

**DOI:** 10.1101/171629

**Authors:** Michael Doron, Giuseppe Chindemi, Eilif Muller, Henry Markram, Idan Segev

## Abstract

The pronounced, long lasting, regenerative NMDA-spike is initiated in individual dendritic branches of different types of neurons and is known to play a key role in dendritic computations and plasticity. Combining dynamic system theory and computational approaches, we systematically analyzed how timed synaptic inhibition activated during the NMDA-spike time-course, sculpts this spike and its associated current influx. When impinging on its early phase, individual GABAergic synapse activation transiently, but strongly, dampened the NMDA-spike; later inhibition prematurely terminated it. This inhibition reduced the NMDA-mediated Ca^2+^ current by up to 60%. NMDA-spikes in distal dendritic branches/spines are longer lasting and more resilient to inhibition, and thus enhance synaptic plasticity at these branches. Examination of this sensitivity of the NMDA-spike to well-timed synaptic inhibition suggests that NMDA-spikes are highly modifiable signals which enable sparse weak distal dendritic inhibition to finely tune both the neuron’s output spikes as well as the branch’s/spine’s Ca^2+^ current associated with the local NMDA spike.

## Introduction

The possibility that the nonlinear properties of the dendritic membrane might endow dendrites with enhanced computational and plastic capabilities was first suggested more than 50 years ago by Rall and colleagues (Rall et al. 1966; Segev & Rall 1988; Rall & Shepherd 1968), and more recently in the influential studies by Mel (Mel 1992; Mel 1993) and Poirazi and Mel (Poirazi & Mel 2001; Poirazi et al. 2003b). Impressive advances in molecular and optical technique have demonstrated that dendrites are particularly rich in a myriad of nonlinear membrane ion channels (Johnston & Narayanan 2008; Magee 2016) and that these channels are involved in specific plastic, computational and cognitive processes (Lavzin et al. 2012; Smith et al. 2013; Poirazi & Mel 2001; Poirazi et al. 2003a; Palmer et al. 2014; Hay & Segev 2015; Hoffman et al. 1997; Golding et al. 2002; Major et al. 2013; Cuntz et al. 2014). The recently discovered Na^+^-, Ca^2+^- and NMDA-dendritic spikes, (Major et al. 2013; Takahashi et al. 2016; Larkum et al. 1999; Schiller et al. 2000; Golding & Spruston 1998) are of particular interest. One of the most significant is the long-lasting NMDA spike (10-100 milliseconds in duration), which was shown to be generated locally in multiple apical and basal dendritic braches of both cortical and hippocampal pyramidal neurons (Poleg-Polsky 2015; Branco & Häusser 2011; Milojkovic et al. 2004; Nevian et al. 2007; Lavzin et al. 2012; Major et al. 2013; Schiller et al. 2000). It has been shown that dendritic Na^+^-, Ca^2+^- and NMDA-spikes can implement a variety of computational functions, including input pattern classifications (Mel 1992), coincidence detection (Larkum & Nevian 2008; Schiller & Schiller 2001), and directional selectivity (Smith et al. 2013; Branco et al. 2011). Importantly, the Ca^2+^ influx associated with dendritic spikes plays a key role in modulating the plasticity of dendritic synapses (Gordon et al. 2006; Sandler et al. 2016; Gambino et al. 2014; Villa et al. 2016).

Nonlinear dendritic signals could be effectively modulated by dendritic inhibition. Theoretical studies have shown that a well-located dendritic inhibition, when preceding its respective dendritic/somatic spikes, could either completely abolish these spikes or modulate their amplitude (Rhodes 2006; Gidon & Segev 2012; Jadi et al. 2012). Experimentally, it was recently demonstrated that NMDA spikes, and their resulting somatic bursts of Na^+^ spikes, are regulated by SOM^+^ interneurons (Lovett-Barron et al. 2012). A recent study (Müllner et al. 2015) showed that a single GABAergic contact could strongly, and very locally, reduce the influx of Ca^2+^ current originating from the back propagating action potential. As a whole, these studies convincingly showed that sparse dendritic inhibition could effectively modulate dendritic excitability and its associated plasticity inducing signals (Bar-Ilan et al. 2013). It is worth noting in this context that in both the cerebral cortex and the hippocampus, different classes of inhibitory interneurons target pyramidal neuron dendrites in different dendritic domains (Stokes et al. 2014; Markram et al. 2004; Fishell & Tamás 2014; Bloss et al. 2016) and are activated in different behavioral states such as sleep cycles (Ma et al. 2010; Pouille 2001; Pouille et al. 2009; Wehr & Zador 2003). This raises the interesting possibility that the various types of interneurons might selectively control different nonlinear dendritic signals. For instance, Martinotti cells that mostly target the distal apical tree might selectively control NMDA spikes in distal apical dendrites, whereas basket inhibition might selectively control the Ca^2+^ generated at the main apical branch of L5 and L2/3 pyramidal cells, whereas chandelier inhibition (targeting the initial segment of the axon of pyramidal neurons) could selectively control the somatic/axonal Na^+^ spike (see discussion by (Gidon & Segev 2012)).

But how can synaptic inhibition interact with the NMDA spike after it has already been initiated? Because the NMDA spike is long lasting (Major et al. 2008), it is likely that synaptic activity *in vivo* will bombard the NMDA during its plateau phase. This is particularly true for synaptic inhibition, which is activated *in vivo* at a relatively high frequency after excitation (Ma et al. 2010; Pouille 2001; Pouille et al. 2009; Wehr & Zador 2003). How would the NMDA spike “react” to such well-timed inhibition? Would it typically be completely abolished or should we expect that the NMDA spike, and its associated current influx, be finely tuned by such inhibition? We explored these questions by using the most advanced model of the NMDA spike, realistic morphology and the properties of inhibitory synapses. We applied dynamic system theory combined with computer simulations, to explore how inhibition interacts with and affects the dynamics of the NMDA spike. Employing detailed models of a 3D reconstructed L5 cortical pyramidal cell, we found that an early activated single GABAergic synapse “riding” on the plateau phase of the NMDA spike could significantly, but transiently, dampen that spike. At later arrival times, the same weak inhibition fully terminated the NMDA spike prematurely. We also found that in distal apical tuft dendrites, NMDA spikes are more resistant to synaptic inhibition than NMDA spikes in proximal dendrites. These findings suggest that due to the slow kinetics and unique membrane mechanisms underlying the NMDA spike, this spike is more susceptible to being sensitively modulated by well-timed local dendritic inhibition. This inhibition might gradually tune both the global spiking output of the neuron as well as the plasticity of synapses at the level of individual dendritic branches and their respective dendritic spines.

## Results

### Sculpting the NMDA spike voltage trajectory by timed synaptic inhibition

Figure 1 illustrates the strong impact that timed weak dendritic inhibition can have on the NMDA spike. The modeled NMDA spike was initiated by simultaneously activating 20 excitatory AMPA- and NMDA-based synapses (0.4 nS peak conductance, red synapses in Figure 1A, see Methods) impinging on a single distal apical branch of a 3D reconstructed layer 5 neocortical pyramidal cell. A single GABA_A_ mediated inhibitory synapse (1 nS peak conductance, blue synapse in Figure 1A, see Methods) located in the center of the excitatory synapses was activated at various times relative to the activation of the excitatory synapses. As expected, this rather small inhibitory conductance had a very small effect on the voltage trajectory of the NMDA spike when it preceded excitation (Inhibition arriving either 10 msec before excitation or right at the onset of excitation, Figure 1B dark red and green NMDA spikes, respectively). The same weak inhibition generated stronger hyperpolarization when activated at the plateau phase of the NMDA spike, at a delay of Δt = 10 msec after excitation. Surprisingly, the NMDA spike fully recovered from this strong hyperpolarization, and returned the voltage to its original trajectory following inhibition (Figure 1B, blue trace). More surprising still, when inhibition was further delayed (Δt = 20 msec), the NMDA spike was prematurely terminated (Figure 1B, lower right), and the voltage continued to hyperpolarize after the inhibitory input.

**Figure 1.**
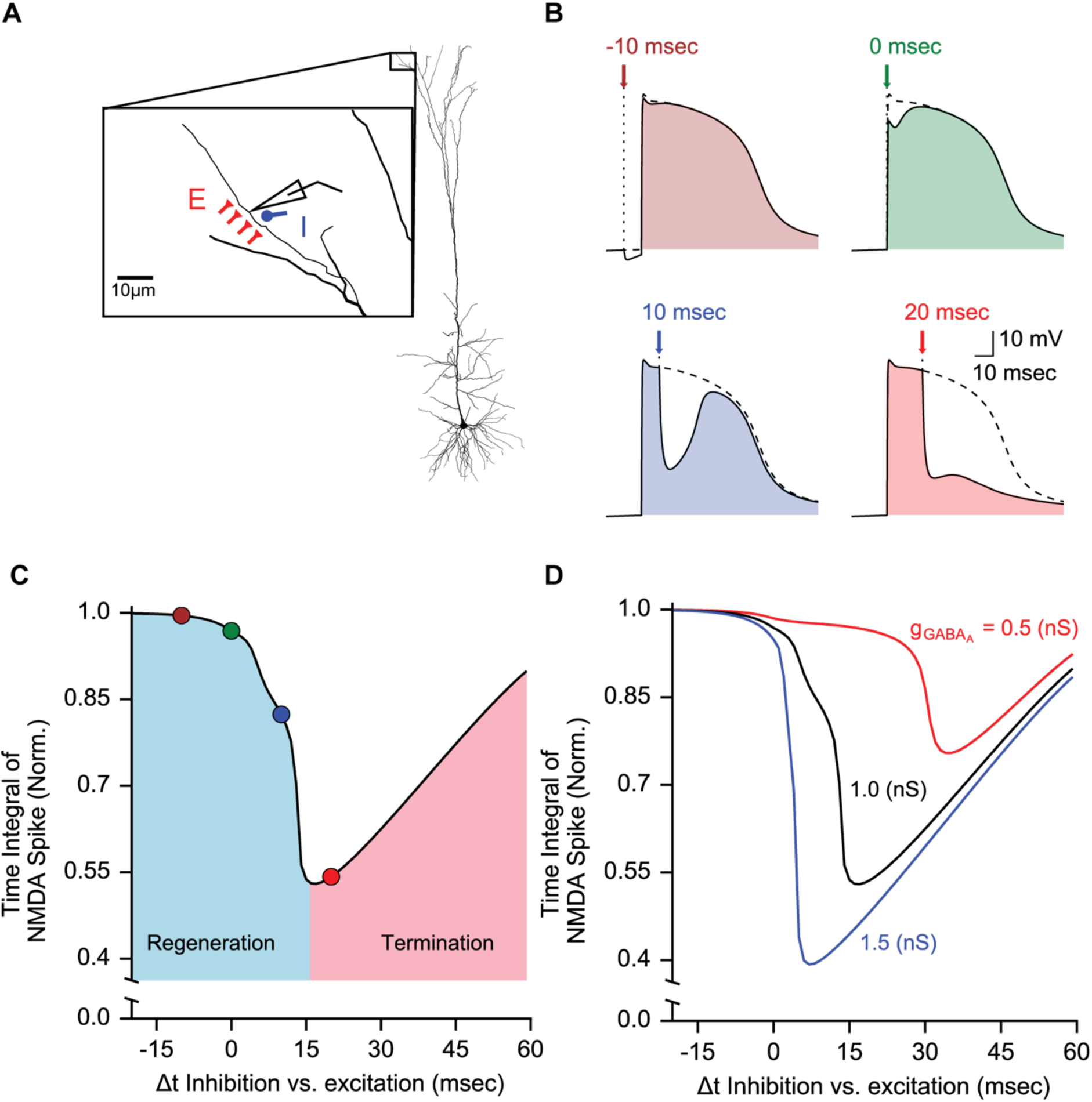
Dendritic NMDA spikes are particularly susceptible to synaptic inhibition. (A) Reconstructed L5 pyramidal neuron model used in the present study. Twenty synchronously activated AMPA- and NMDA-based excitatory synapses (0.4 nS peak conductance each, schematic red synapses) were distributed around the center of a distal apical dendritic branch, with 1 synapse per 2.2 μm. A single GABA_A_ synapse (blue synapses, 1 nS peak conductance) was placed in the center of the excitatory synapses from which the membrane voltage was recorded (schematic electrode). (B) NMDA spikes generated in the modeled dendritic branch shown in (A); the same inhibitory synapse was activated at four different times with respect to the activation of the excitatory synapses (the arrows point to the inhibition activation time). The colored areas show the NMDA spike in the presence of inhibition as compared to the control trace without inhibition (black trace). (C) Normalized time integral of the NMDA spike for different activation times of the inhibitory synapse (the “vulnerability function”, see Results). Each colored dot on the black curve corresponds to the respective traces in (B). The regeneration phase denotes the cases in which the NMDA spike fully recovered to its original trajectory following inhibition; the termination phase indicates the instances in which the NMDA spike was terminated prematurely following inhibition. (D). Same as (C), with different GABA_A_ peak conductance values as indicated. The larger the GABA_A_ conductance, the larger the drop in the NMDA time integral and the sooner the regeneration phase ends and the Termination phase starts.

In order to quantify this sensitivity of the NMDA spike to timed inhibition “riding” on its plateau potential, we plotted the time integrals of the NMDA spike normalized to control conditions (without inhibition, dashed traces in Figure 1B) as a function of Δt - the activation time of inhibition relative to excitation (Figure 1C). We term this the “vulnerability function” of the NMDA spike to inhibition. The effect of inhibition during the time course of the NMDA spike could be divided into two distinct regimes, which we call the “regeneration” and “termination” phases. In the “regeneration” phase, the NMDA plateau hyperpolarizes due to the inhibition and then fully recovers to its original trajectory at the succession of inhibition (light blue region in Figure 1C). In this regenerative phase, the time integral of the NMDA spike voltage dropped (by 1 nS GABA_A_ inhibition) continuously to up to ~45% of its original time integral (at about Δt = 15 msec for the case shown in Figure 1). For larger Δt values, the NMDA spike was prematurely terminated by the inhibition (red region in Figure 1C and see the corresponding example in Figure 1B for Δt = 20 msec). After the NMDA spike entered the “termination” stage, its time integral increased almost linearly when the inhibition was activated at progressively later times. For larger GABAergic conductance, the termination phase started earlier and the time integral of the NMDA spike was further reduced by inhibition (Figure 1D). This two-regime sensitivity of the NMDA spike to timed inhibition was robust across different parameter ranges and across different models of the NMDA receptor current, (Figures S1 and S2). Interestingly, the sensitivity of the dendritic Ca^2+^-spike to timed inhibition was qualitatively different than that of the NMDA spike, as the NMDA spike is easier to terminate with a delayed inhibition, while the Ca^2+^ spike is easier to terminate in its onset, while being more robust later on (figure S4, also see (Müllner et al. 2015)).

### Distal dendritic inhibition is particularly effective in modulating the NMDA spike

We next examined how the spatial location and activation timing of the inhibitory synapse affect the NMDA spike. In Figure 2, the same configuration as in Figure 1A was used but the location of the inhibitory the GABA_A_ synapse was shifted with respect to the location of the excitatory synapses (Figure 2A). The inhibitory synapse was placed either at 35 *μ* m distal to the center of the dendritic branch, in its center, or 35 *μ* m proximal to it (Figure 2A). The weak (1 nS peak) inhibitory synapse, when located 35 *μ* m distal to the NMDA spike and activated at Δt of 20 msec, terminated the NMDA spike (red NMDA spike in Figure 2B). The same inhibitory synapse hardly affected the NMDA spike when placed proximally to it (blue NMDA spike in Figure 2B). Furthermore, the timing of the inhibition that maximally affected the time integral of the NMDA spike was earlier for the distal synapse and later (and less effective) for the proximal synapse (Figure 2C, red versus blue spikes, respectively). When comparing the vulnerability function of the NMDA spike across the different spatial locations of the inhibition, this function deepened significantly when the inhibitory synapse was located either directly at (x = 0) or distally to the NMDA spike, at x = +35 *μ* m (Figure 2D). Thus the modulation of the NMDA spike by a branch specific synaptic inhibition appears to be highly effective when the inhibition is located either “on spot” or distally to the NMDA spike initiation site. This finding is consistent with (Gidon & Segev 2012) who studied the impact of dendritic inhibition on the more global dendritic Ca^2+^ spike generated at the main branch of the apical dendrite.

**Figure 2.**
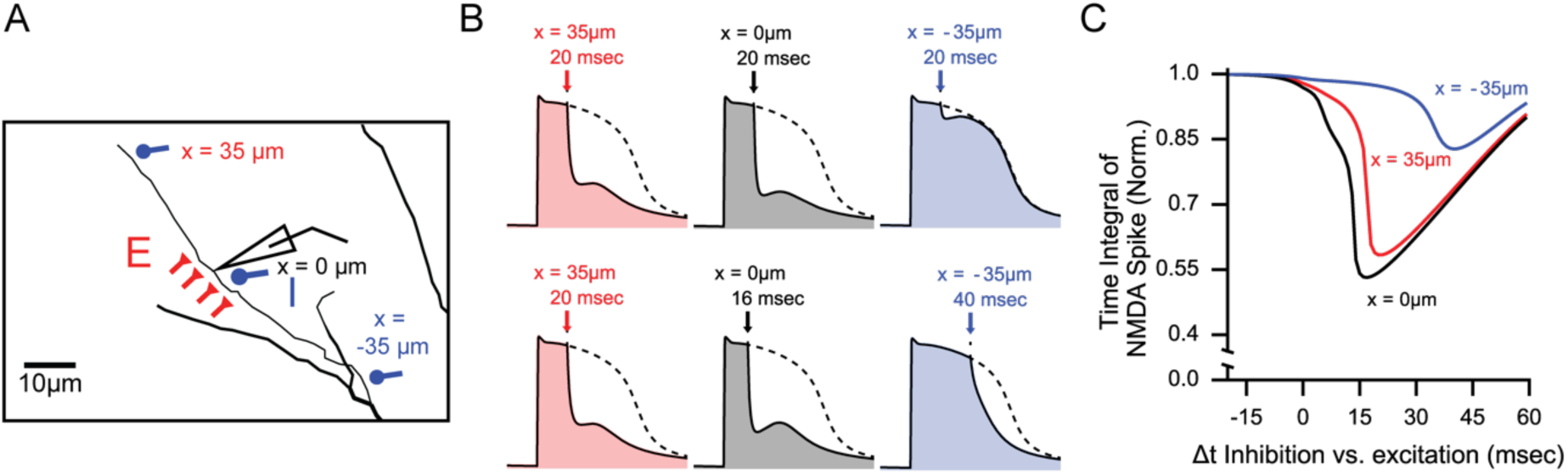
NMDA spikes are affected more by distal than proximal inhibitory synapses. (A) Similar to Figure 1A, with three locations, x, of the inhibitory synapse with respect to the center of the modeled dendritic branch where the excitatory synapses are located. For x = 0, inhibition impinges in the middle of the branch; for 35 μm it is distal to the soma and for -35 μm it is proximal to the soma. (B) (Top) Similar to Figure 1B with inhibition arriving 20 milliseconds after excitation. Each trace shows the NMDA spike for the three inhibitory locations as in (A); with “on spot” inhibition (x = 0, black), dampening the NMDA maximally, distal inhibition is almost as effective (red), whereas the proximal inhibition hardly affects the NMDA spike (blue). (Bottom) Same as top, with the inhibition arriving at the time where for each location, it reduced the NMDA maximally. (C) Similar to Figure 1D, where each curve depicts the effect of inhibition when placed at different dendritic locations and activated at different times.

### Dynamic analysis and mechanistic explanation of the bi-phasic effect of inhibition impinging on the NMDA spike

We studied the behavior of the V-I relationship of the NMDA spike in the presence of synaptic inhibition as a dynamical system, using a single compartment neuron model consisting of leak ion channels and AMPA-, NMDA- and GABA_A_-based synapses (peak conductance of 7.5 nS, 7.5 nS and 1.5 nS respectively, see Methods). As was shown by (Schiller & Schiller 2001; Major et al. 2013; Jadi et al. 2012; Sanders et al. 2013), the NMDA current (red trace in Figure 3A) when accompanied by the leak current (blue trace in Figure 3A with reversed sign) creates a bi-stable dynamical system with three fixed points (the three circles in Figure 3A). At these fixed points, the outward and inward currents are equal such that the net membrane current is zero. By perturbing the voltage around the fixed points, the left-most intersection (the blue circle labeled “lower stable” in Figure 3A) emerges as a stable fixed point, because a small hyperpolarization from this point will produce an inward current that will result in depolarization back to the fixed point, whereas a small depolarization will produce an outward current that will result in hyperpolarization back to the fixed point (blue and red arrows around blue circle, Figure 3A). Similarly, the right-most intersection (the red circle labeled “upper stable” in Figure 3A) is also a stable fixed point. The middle intersection, however, is an unstable fixed point; a small depolarization will produce an inward current that will further depolarize the membrane and a small hyperpolarization from this point will produce an outward current that will result in further hyperpolarization. This point was thus dubbed the “unstable threshold”; i.e., the critical voltage beyond which due to depolarization, the membrane voltage will change regeneratively, thus initiating the NMDA spike ((Jack et al. 1975), Figure 8.11 and see also (Schiller & Schiller 2001; Major et al. 2013; Jadi et al. 2012)).

**Figure 3.**
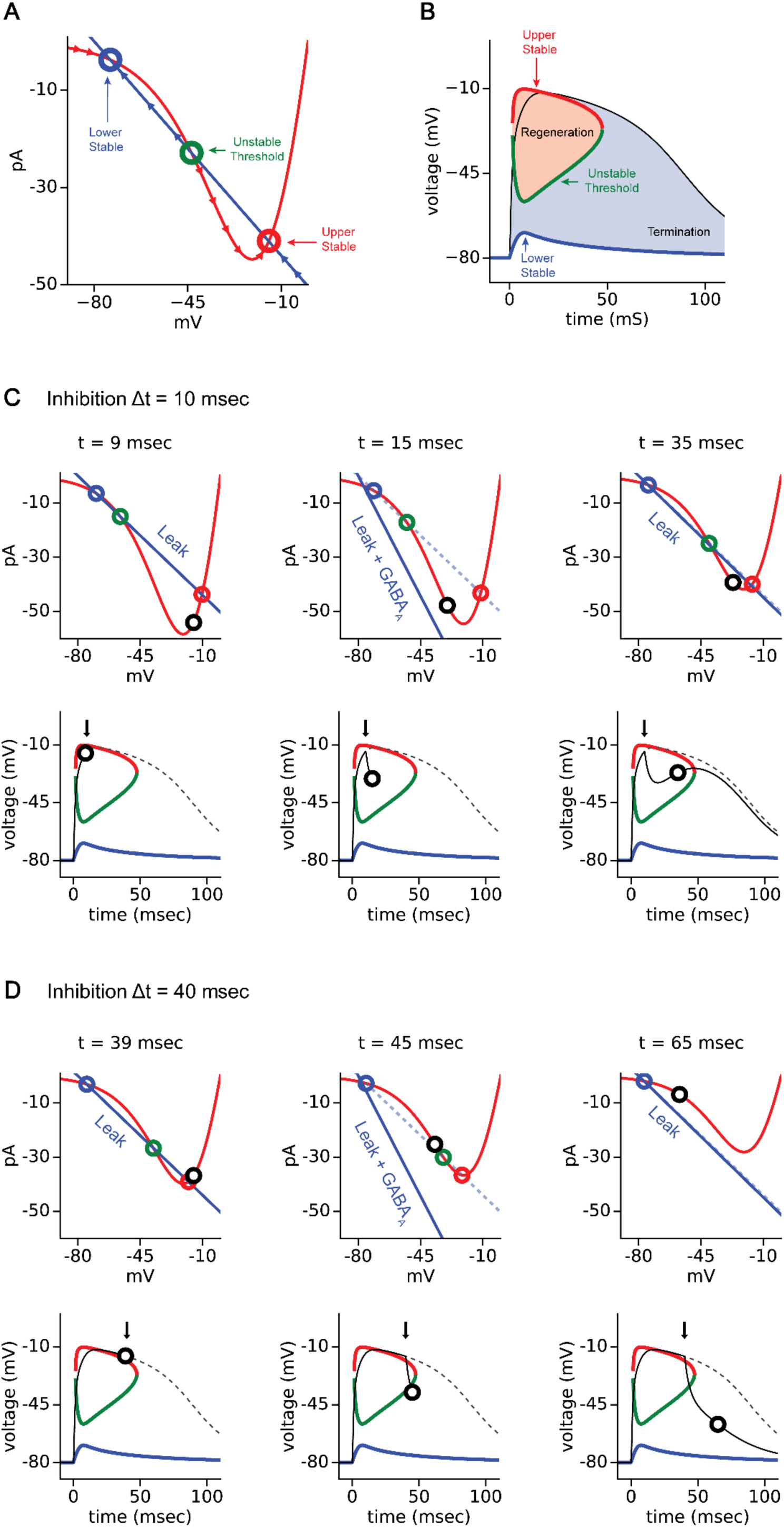
mechanistic explanation of the dynamic vulnerability of the NMDA spike to timed inhibition. (A) The Voltage Current (V-I) curve for the NMDA current (red trace) and for the leak current with reversed sign (blue line) in an isopotential neuron model. These currents were measured at a fixed time of 30 milliseconds following the onset of synaptic excitation (see Methods). The intersections between these two currents are marked by the colored circles, which constitute three fixed points. The middle point (green, “unstable threshold”) is unstable and the other two are stable (blue, “lower stable” and red, “upper stable”). Arrows on the red and blue curves indicate the direction of the voltage at that time point subsequent to a small perturbation around these fixed points (see (Jack et al. 1975) Figs. 8.11 and 8.12, and (Major et al. 2013)). (B) The trajectory of the three fixed points over the time course of the NMDA receptor activation (red, green, blue as in (A)), plotted together with the voltage trajectory of the NMDA spike (black). The red area labeled “regeneration” marks the region where a small hyperpolarization from the upper fixed point (red curve) and a small depolarization from the unstable threshold (green curve) resulted in further depolarization of the voltage. The blue area labeled “termination” marks the region where after a small voltage perturbation the voltage further hyperpolarized. (C) Inhibition (at Δt = 10 msec) during the early phase of the NMDA spike hyperpolarized the voltage but it remained in the regeneration regime, allowing the NMDA spike to fully recover following inhibition. Top: Similar to (A) for three time points (shown above) along the time-course of the NMDA spike. Dashed blue lines depicts the leak current, solid blue lines depict the leak + the inhibitory GABA_A_ current (reversed in sign) at the respective times. The black circle represents the voltage at these time points. Bottom: Similar to (B). The NMDA spike without inhibition is shown by the dashed line; voltage trace of the NMDA spike with inhibition up to that respective time point depicted above is shown by the solid black line in the left and middle frames. The black circle represents the voltage at the respective time points. The black arrow marks the time of activation of the inhibitory synapse. The solid black line in the rightmost frame shows the whole voltage trajectory of the NMDA spike following the inhibition. (D) Same as in (C) but with late inhibition arriving at Δt = 40 msec. In this case inhibition hyperpolarized the NMDA spike to below threshold, thus terminating the NMDA spike.

Because the inward NMDA current changes over time, the voltage values where the inward NMDA current and the outward leak current are equal also change over time. Consequently, the fixed points change during the time course of the NMDA spike (Figure 3B). Since a small hyperpolarization from the upper fixed point will result in further depolarization and a small depolarization from the middle fixed point will also result in further depolarization, we labeled this the “regeneration” region of the NMDA spike (shaded light red region in Figure 3B). The complementary shaded blue region in Figure 3B was termed the “termination” region of the NMDA spike because a small perturbation of the voltage inside it (in both the hyperpolarized and depolarized directions) will result in further hyperpolarization. Examination of the trajectories of the fixed points shows that during the time course of the NMDA spike, the threshold (the green curve in Figure 3B) is relatively hyperpolarized in the early stages, and progressively becomes more depolarized with time. This accounts for the recovery of the NMDA spike subsequent to early inhibition “riding” on its plateau phase and the premature termination of the NMDA spike subsequent to later inhibition as shown in Figure 1B, since inhibition-induced hyperpolarization that does not cross the threshold in its early hyperpolarized state may still cross the threshold when arriving later in its depolarized state.

This behavior is depicted in more detail in Figure 3C which illustrates the case of early inhibition arriving at Δt = 10 msec following the initiation of the NMDA spike. The three V-I curves (top) and the respective voltage trajectories (bottom) were sampled at t = 9, 15 and 35 msec during the NMDA spike. At t = 9 msec (before inhibition was activated), the NMDA spike reached a value of -12 mV (black circle at left top and bottom frames in Figure 3C) and was still depolarizing, as it was in the regeneration region. Following the activation of inhibition (at t = 10 msec), the outward GABA_A_-dependent current was added to the outward leak current, thus shifting the dashed blue curve (the leak current) to the steeper solid blue line (Figure 3C, top middle frame). At t = 15 msec there was only a single (stable) intersection point between the solid blue and the red V-I curves, with the inhibition-induced outward current hyperpolarizing the NMDA spike (Figure 3C, bottom middle frame). This inhibitory current continued to hyperpolarize the NMDA spike transiently, until the GABA_A_ - current ended. At later inhibition arrival times (e.g., At t = 35 msec) the outward current converged back to the leak current (solid blue superimposed on the dashed blue line in Figure 3C, top right). In the case here, this early inhibition hyperpolarized the NMDA spike voltage to -30 mV (black circle in Figure 3C, middle frames), which was still more depolarized than the threshold (green line in Figure 3C, bottom frame) such that the voltage remained in the “regeneration” region, allowing the NMDA spike to recover back to the upper stable fixed point (Figure 3C, bottom right).

In contrast, when the inhibition arrived later, at Δt = 40 msec (Figure 3D), the NMDA spike threshold was more depolarized than the NMDA threshold at earlier times. In this case, for the same inhibitory conductance as in Figure 3C, and because the V-I curve of the NMDA current was shallower at these later times, the hyperpolarization due to inhibition crossed the threshold for the NMDA spike (green circle in middle top and green curve in bottom middle of Figure 3D, respectively). Thus, the voltage perturbation due to inhibition resided in the “termination” region that further hyperpolarized the NMDA spike until its premature termination.

### Branch-specific NMDA spike “vulnerability function”

The simulations so far have focused on a single distal apical branch in a passive model of an L5 pyramidal cell ((Hay et al. 2011), without active channels). In order to further characterize the influence of timed inhibition on NMDA spikes and examine its robustness over different dendritic branches, we ran the same simulations on all terminal dendrites of the cell as well as on a single compartment model with various passive properties (Figure S5).

For a fixed NMDA/AMPA conductance, the NMDA spikes in dendritic branches with large input resistance were longer lasting and more resilient to GABA_A_ inhibition as compared to branches with small input resistance. This is illustrated in the two examples in Figure 4A, where a long lasting (61 msec) NMDA spike was generated in a distal apical branch with large input resistance (1285 MΩ) whereas a much briefer NMDA spike (25 msec) was generated in an oblique branch with a smaller input resistance (810 MΩ). The former NMDA spike recovered after GABAergic inhibition at *Δt* = 10 msec whereas the latter NMDA spike terminated after the same inhibition, suggesting that the NMDA spikes in branches with a larger input resistance are longer lasting and more resilient to inhibition (Figure 4B). We further computed the Pearson correlation coefficient between various passive properties of the branch (its diameter, length, surface area, distance from the soma and input resistance) and the critical GABA_A_-conductance required to terminate the NMDA spike (Figure 4C). This critical GABA_A_ conductance was strongly correlated with the input conductance of the dendritic branch; the larger the input resistance (the smaller the input conductance), the larger the GABA_A_ conductance required to terminate the NMDA spike. The critical GABA_A_ conductance was also correlated (but less strongly) with the distance of the dendritic branch from the soma (Figure 4C). These results, together with those reported in (Poleg-Polsky 2015; Major et al. 2008) showing that branches with larger input resistance are more prone to generate an NMDA spike, imply that NMDA spike based plasticity may be more likely to take place at distal branches with large input resistance.

**Figure 4.**
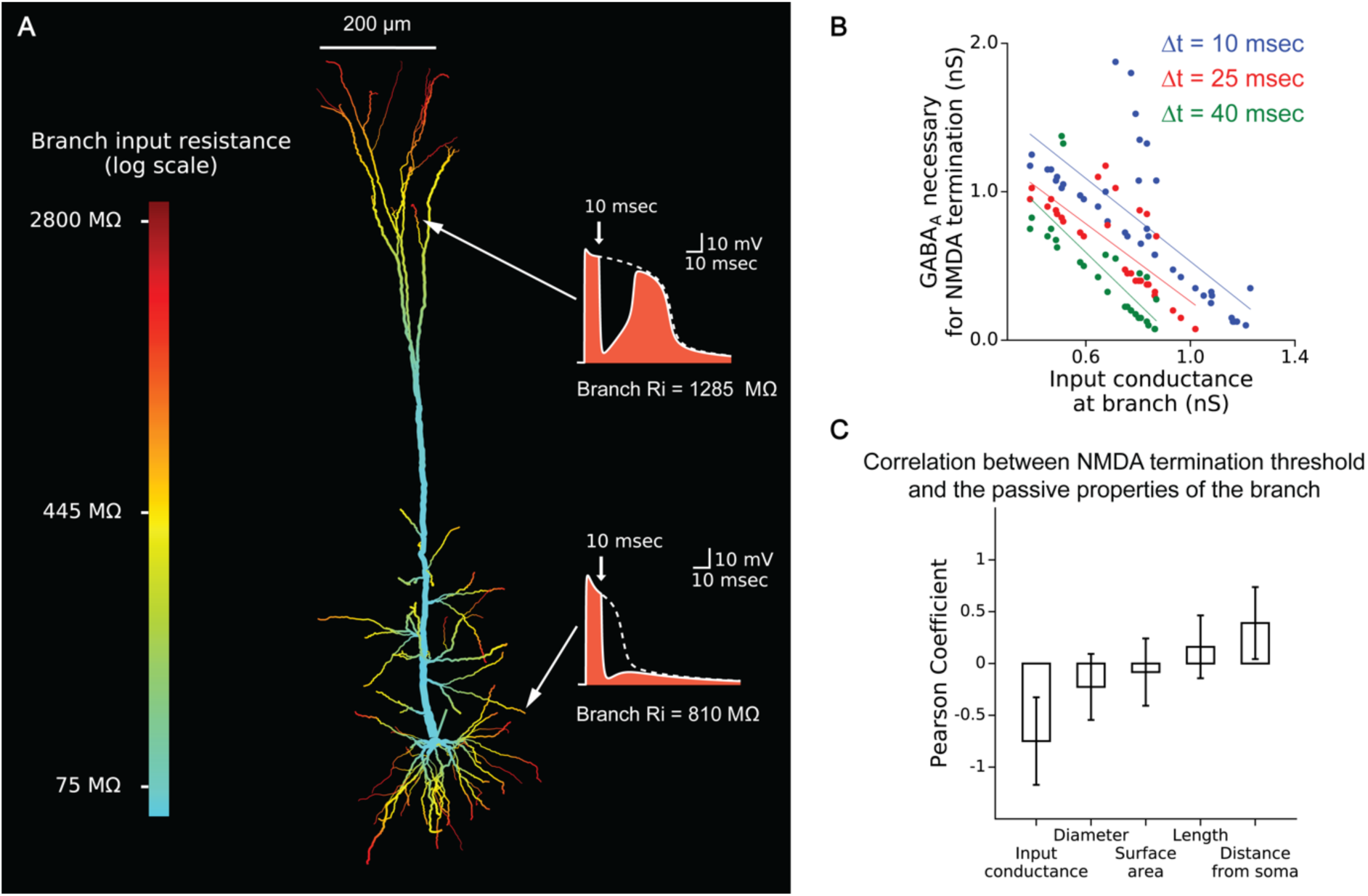
NMDA spikes in dendritic branches with large input resistance are more resilient to timed inhibition. (A) The NMDA spike in dendritic branches with large input resistance (top right) is long lasting and more resilient to timed inhibition than the NMDA spike in dendritic branches with a smaller input resistance (lower trace at right). The white dashed traces show the NMDA spikes in control conditions; the NMDA spike with inhibition (8 nS peak conductance) activated at Δt = 10 msec (red arrows) is shown in white. The NMDA spikes were generated as in Figure 1; excitatory synapses were distributed uniformly in the most distal 20 μm of the respective branches. White arrows point to the activated branches. The branches of the modeled neuron are colored according to their input resistance (log scale on left). (B) The GABA_A_-conductance required to terminate an NMDA spike was reciprocally and linearly related to the input conductance in the branch. Colored dots depict the GABA_A_-conductance required to terminate NMDA spikes at the various branches, with 3 cases of inhibition arriving at different delays, Δt, with respect to excitation. (C) Correlation between the critical GABA_A_ conductance required to terminate the NMDA spike and various properties of the dendritic branches. Note that the input conductance of the branch was strongly correlated with the GABA_A_ conductance needed to terminate the NMDA spike. Other passive properties (branch diameter, surface area, length) were not correlated with the critical GABA_A_ conductance; the distance of the branch from the soma was only partially correlated with the critical GABA_A_ conductance. In this figure, the Hay (Hay et al. 2011) model was used, without active channels.

Analysis of the V-I curve of the NMDA spike in modeled cells with different input resistances (Figure S5A) helped explain the resilience of the NMDA spike to inhibition when the input resistance is large in two complementing ways. The initiation threshold of the NMDA spike was more depolarized in branches with low input resistance. This means that it was easier for the inhibition-induced hyperpolarization to cross this threshold and, thus, prematurely terminate the NMDA spike. In addition, the upper fixed point (the plateau voltage) was more hyperpolarized in branches with low input resistance. Hence, it was easier for the inhibition impinging on this smaller plateau to cross the NMDA spike threshold and terminate this spike. This increased effect of inhibition is counter intuitive to the notion that lower input resistance always decreases the inhibitory hyperpolarization, since the relative hyperpolarization is indeed lower in lower input resistance cases. However, as can be seen in Figure S5A, the decreased input resistance manifests as an increase in leak current, thus resulting in a larger total inhibitory current, thus increasing the vulnerability of the NMDA spike – larger total inhibitory current means higher NMDA threshold, lower NMDA plateau voltage, and less inhibitory conductance necessary for termination of the NMDA spike. This can be seen in Figure S5B,C, where two NMDA spikes created with the same NMDA conductance at higher and lower input resistance conditions responded differently to the inhibition of a single GABA_A_ synapse (1 nS) arriving 10 milliseconds after excitation. By plotting the inhibitory conductance required to terminate the NMDA spike, we found that for a given input resistance and time interval, the critical GABA_A_ conductance increased linearly with the NMDA conductance. As the input resistance decreased, the same GABA_A_ conductance terminated the NMDA spike when arriving at shorter delays following excitation.

Thus the vulnerability of the NMDA spike to timed inhibition appears to depend strongly on the cable properties of the dendritic branch where this spike is generated. Distal dendritic branches with a large input resistance are more prone to generating an NMDA spike to start with (Polsky, 2015). When generated, this spike is long lasting (on the range of ~100 msec) at these branches and is more resilient to inhibition as compared to the briefer NMDA spikes that are generated at dendritic branches with lower input resistance.

### Timed inhibition shifts the duration distribution of the NMDA spike population from unimodal to bimodal

Given the finding here that the NMDA spikes have different durations at different dendritic branches and that these spikes can either fully recover or terminate following timed inhibition, we tested in what ways the duration distribution of the dendritic NMDA spikes population changed with inhibition. Using the full nonlinear L5 neuron model of (Hay et al. 2011) we found that in the absence of inhibition, the distribution of the duration of the dendritic NMDA spikes was unimodal, with some spikes lasting only ~30 msec but others that could last up to 150 msec in the parameter range we used (Figure 5A). Timed inhibition polarized this distribution into two distinct populations of NMDA spikes: the spikes that were terminated (becoming as short as 10 msec in duration) and those that were recovered following inhibition, and remained long lasting after inhibition (Figure 5B-D). The difference between these two populations was significant both in terms of the duration of the NMDA spikes before and after inhibition (Figure 5E), and as regards to the time integral of the respective NMDA based current (Figure 5F). The pattern of timed inhibition terminating the weaker NMDA spikes and keeping the most resilient ones, with later inhibition terminating a larger portion of the NMDA spikes, can be thought of as a mechanism for controlling local plasticity in the dendrites by allowing only the strongest local dendritic spikes to survive and influence the cell’s plasticity (see Discussion).

**Figure 5.**
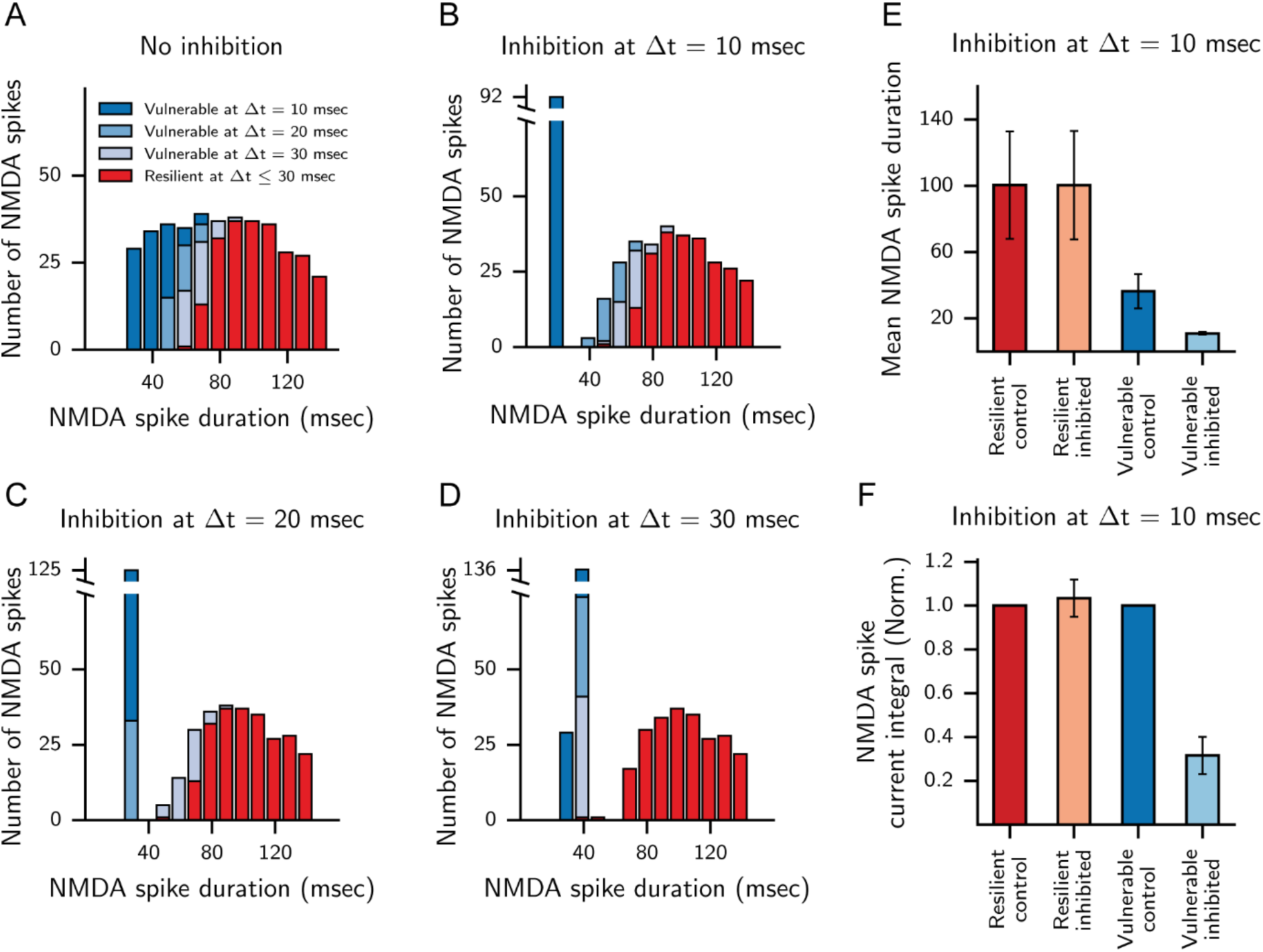
Timed dendritic inhibition shifts the duration distribution of the NMDA spikes population from unimodal to bimodal. (A) The unimodal distribution of NMDA spike durations in the absence of inhibition. Bars are the histogram of the duration of NMDA spikes in different dendritic terminals of the modeled cell shown in Figure 1. In these simulations, different NMDA levels of conductance were used along with a single inhibitory conductance of 1 nS (see Methods). Colors represent the NMDA spikes that were terminated with inhibition arriving at different times as shown above. (B-D). Progressively delayed inhibition terminated the shorter NMDA spikes (different blue colors), leaving the longer NMDA spikes relatively unchanged (red spikes). (E) The duration of the vulnerable NMDA spikes significantly decreased following inhibition (blue), whereas the duration of the resilient NMDA spikes remained unchanged (red). (F) As in (E) but for the normalized current time integral. In this figure, the full nonlinear model in (Hay et al. 2011) was used.

### NMDA spikes generated in multiple spines: Effects of dendritic versus spine inhibition

Recent work has shown that the effect of inhibition is highly localized when synaptic inhibition impinges on the dendritic spine together with an excitatory synapse located on the same spine (Chiu et al. 2013; Higley 2014; Higley & Sabatini 2012). How would spine inhibition interact with the dendritic NMDA spike when generated by a group of excitatory spine synapses, compared to dendritic inhibition? To examine this question, we used the spine model in (Grunditz et al. 2008) and distributed 20 such spines on the distal 20 *μ*m of an apical branch of a CA1 modeled pyramidal cell. A single GABA_A_ synapse was located either on the dendrite itself or on a single spine head. Figure 6A shows that when placed on the dendritic branch (as is typically the case), timed inhibition could terminate the NMDA spike at all activated spines (for Δt = 20 msec, red trace in Figure 6B). When the same inhibitory synapse was placed at the spine head (Figure 6D), the NMDA spike was not terminated at any of the activated spines (Figure 6E). This was true despite the observation that inhibition on the spine head hyperpolarized that specific spine voltage more than inhibition on the dendrite (Compare Figure 6B left and Figure 6E left). In other words, the cooperative NMDA spike, generated collaboratively at all 20 activated spines, protected the NMDA spikes at individual spines from local inhibitory input. Furthermore, the concentration of Ca^2+^ flowing into the spine head through the NMDA receptors dropped by ~50% when inhibition impinged on the dendrite (Figure 6C). The same inhibition when located on the spine head reduced the total Ca^2+^ concentration at that spine by only 20% (Figure 6F). The vulnerability function for both the NMDA voltage time-integral (Figure 6G) and for the time integral of the total Ca2+ concentration (Figure 6H) demonstrate the increased effect of dendritic versus spine inhibition. Because intracellular Ca^2+^ concentration is critically involved in plastic processes, the results of Figure 6 have important implications for branch-specific plasticity processes as elaborated on in the Discussion.

**Figure 6.**
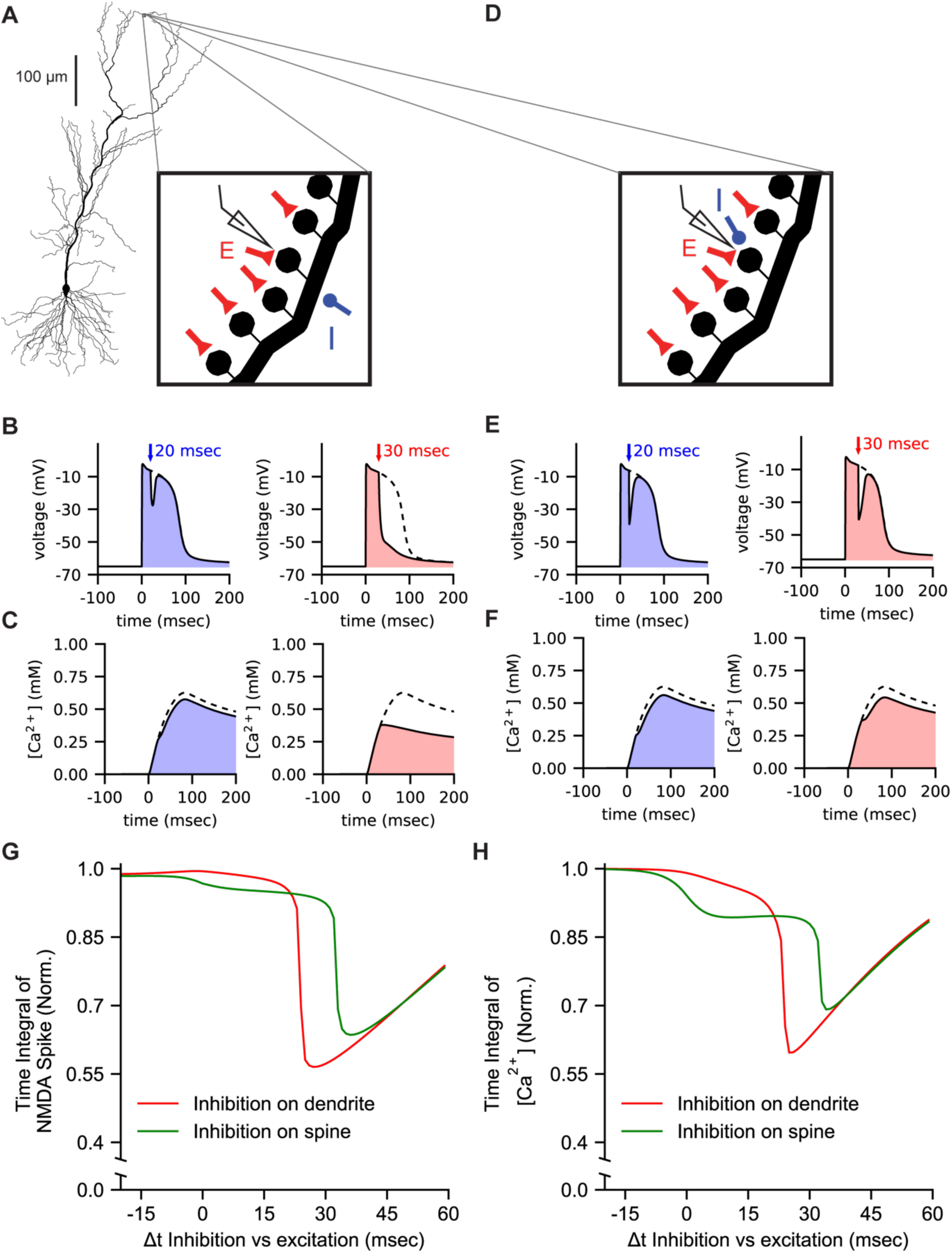
When generated synchronously in multiple spines, the NMDA spike and its corresponding Ca^2+^ influx are strongly affected by dendritic inhibition and are less affected by spine inhibition. (A) CA1 pyramidal neuron model as in (Grunditz et al. 2008). Twenty spines were distributed on the most distal 20 μm of a distal apical branch, where each spine head contained AMPA- and NMDA-based excitatory synapses (0.8 nS peak conductance each, schematic red synapses). A single GABA_A_ synapse (blue synapses, 0.5 nS peak conductance) was placed on the dendrite, in the center of the modeled spines. Voltage and the Ca^2+^ concentration were recorded from the central spine (schematic electrode). (B) NMDA spikes generated by activating the excitatory synapses synchronously on all 20 modeled spines in the modeled dendritic branch shown in (A). The same inhibitory synapse was activated at two different times with respect to the activation of the excitatory synapses (Δt = 10 msec and 20 msec). The colored areas show the NMDA spike in the presence of inhibition as compared to the control trace without inhibition (black dashed trace). (C) Total [Ca^2+^] in the spine head during the NMDA spike without inhibition (black dashed traces) and with inhibition arriving at different times (colored solid traces). (D) Similar to (A), with inhibition impinging on the spine head. (E) Similar to (B). Inhibition arriving on the spine head hyperpolarized the voltage at that spine more than when the inhibition was on the dendritic branch (compare the blue trace on the left to the corresponding trace in (B)), but it does not terminate the NMDA spike at that spine (compare red traces in (E) and (B)). (F) Similar to (C). Inhibition in the dendrite reduced the [Ca^2+^] at the spine head significantly more than the inhibition impinging directly on the spine head (compare (F) to (C)). (G) Vulnerability function of the NMDA spike voltage recorded from a single spine when inhibition is located at the spine head (green, as in case D) or at the dendrite (red, as in case A). (H) Vulnerability function of the total calcium concentration [Ca^2+^] in a single spine when inhibition is located at the spine head (green) or at the dendrite (red). In all case the spine neck resistance was 1.2 GΩ.

As shown in Figure 6, for spines with large neck resistance, the local inhibitory shunt at the spine head membrane does not propagate well to the adjacent spines (see also (Gidon & Segev 2012)). This implies that the collective NMDA current generated by the (say 19) adjacent dendritic spines is only slightly affected by this inhibition on a single spine. However, this collective NMDA-mediated current effectively flows from the dendrite into the spine head membrane (in this direction of current flow, the spine base and the spine head are essentially isopotential, (Segev & Rall 1988)). In contrast, when the inhibition is located directly on the stem dendrite, it shunts more effectively the spine head membrane, thus reducing more powerfully the NMDA-current that is generated in each of the activated spines. The above effect diminishes when the spine neck resistance is relatively low and when dendritic input resistance is low (in this case spine inhibition and dendritic inhibition are essentially similar in their effect, see Figure S6).

### Fine modulation of the neuron’s spike output by timed inhibition interacting with dendritic NMDA spikes

We next utilized the full nonlinear model of the L5 pyramidal cell used above (Hay et al. 2011) to examine how branch-specific timed inhibition interacting locally with the NMDA spike controls the output of this cell. Sixteen random dendritic branches were synchronously activated, each by 20 excitatory AMPA and NMDA synapses. In each of these branches two inhibitory GABA_A_ synapses (1 nS each) were also activated at a various Δt, msec with respect to the excitation (Figure 7A). Figure 7C shows the fine modulation of the neuronal output of Na^+^-spikes by dendritic inhibition. In the control condition (no inhibition) the neuron fired a burst of 2.5 spikes on average in response to the 16 activated dendritic NMDA spikes (somatic bursting due to NMDA activation can also be seen in (Polsky et al. 2009; Milojkovic et al. 2004)). When inhibition was activated simultaneously with excitation (second trace from left in Figure 7B, C), only two Na^+^ spikes were generated at the modeled axon, with a further decrease to one spike when Δt = 10 msec and no output spikes with Δt = 20 msec (Figure 7C). When averaging over 60 possible combinations of the 16 activated terminal basal dendritic branches, the average number of spikes evoked in the soma as a response to excitation and inhibition (Figure 7D) resembled the vulnerability function of a single NMDA spike in a single branch (Figure 1C). As a whole, Figure 7 demonstrates that sparse and weak inhibition in distal dendritic braches was able to strongly modulate the neuron’s output when it was well-timed during the plateau phase of local dendritic NMDA spikes.

**Figure 7.**
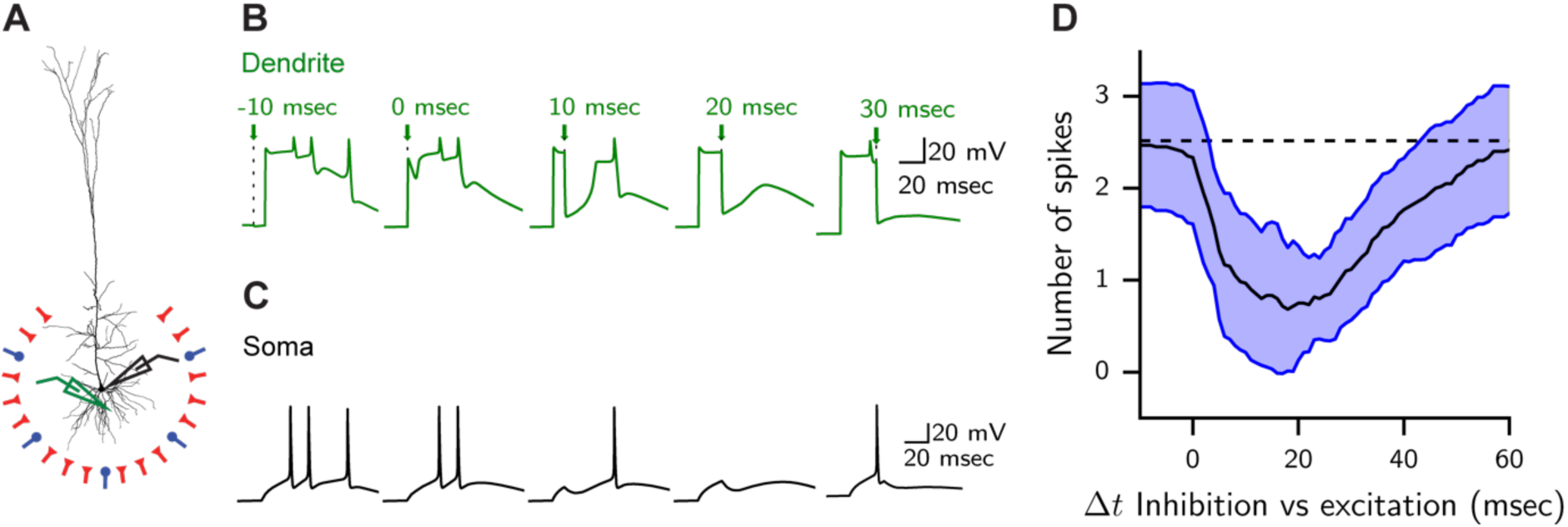
Timed dendritic inhibition interacting with dendritic NMDA spikes finely modulates the neuron’s output. (A) Sixteen random dendritic terminal branches were synchronously activated, each by 20 excitatory AMPA and NMDA synapses (0.4 nS each) and 2 inhibitory GABA_A_ synapses (1 nS each). The excitatory synapses were distributed uniformly over the most distal 20 *μ* m of the activated branch, and the inhibitory synapses were placed 10 *μ* m proximal to the branch termination. All excitatory synapses were activated simultaneously at t = 0 msec and all inhibitory synapses were activated simultaneously at a given Δt msec with respect to excitation. Voltage was recorded from the soma (black electrode) and from one of the activated branches (green electrode). (B) Local NMDA spike modulated by local timed inhibition. Arrows indicate the timing of inhibition with respect to excitation. Recorded from one of the activated branches (green electrode in (A). (C) Soma spikes for the respective cases shown in (B). For early inhibition (Δt = -10 msec) a burst of 3 spikes was generated at the soma. This burst of spikes was finely modulated by weak inhibition impinging at various times during the plateau phase of the dendritic NMDA spikes. (D) Average (black line) and standard deviation (blue lines) of 60 possible combinations of 16 activated terminal basal branches as in (C). The dashed line is the average number of spikes in the control condition without inhibition. In this Figure, the full nonlinear model of (Hay et al. 2011) was used.

## Discussion

Our dynamic system analysis and computational study suggests that due to its unique dynamics, the NMDA spike is particularly sensitive to well-timed and strategically located dendritic inhibition. Inhibition located at the dendritic site of the NMDA spike generation or a few tens of *μ*m distal to it was able to finely and powerfully tune the time course of the NMDA spike and its corresponding current influx (Figures 1 and 2). We showed that there are two distinct regimes for the effect of synaptic inhibition when activated during the long plateau phase of the NMDA spike. In the early phase, weak synaptic inhibition transiently hyperpolarized the NMDA spike; it then fully recovered back to its original trajectory. At later arrival times, at the termination phase, inhibition prematurely terminated the NMDA spike (Figure 1C), decreasing its half width, and not allowing voltage to depolarize back. We showed that this effect remains the same for more distal inhibition, but diminishes significantly for more proximal synapses, making the effect of inhibition on the NMDA spike a branch specific phenomena. We later used the theory of dynamical systems to explain this sensitivity of the NMDA spike to timed inhibition (Figures 3). Depending on the time of inhibition with respect to plateau phase of the NMDA-spike, this inhibition may gradually reduce the NMDA-related Ca^2+^ influx by up to 60% (Figure 6C).

We also explored and mechanistically accounted for the finding that at a certain time-window, synaptic inhibition increases, rather than decreases, the NMDA-current influx (Figure S3) and characterizes the shift the NMDA spike duration over the different dendritic branches from unimodal to bi-modal distribution when a distributed dendritic inhibition is activated (Figure 5). We showed that stronger and longer-lasting NMDA spikes (in distal dendrites with large input resistance) persist in the face of local inhibition, whereas weaker NMDA-spikes (on proximal branches with smaller input resistance) are more susceptible to inhibition (Figures 4 and S5). We next simulated the case of an NMDA spike generated cooperatively in the head membrane of a group of dendritic spines following the activation of their respective excitatory synapses, and showed that timed-inhibition impinging directly on the dendrite effectively controls the NMDA spike in all spines involved, but that the same NMDA spike was resilient to local inhibition on individual dendritic spines (Figure 6). Finally, we demonstrated that weak inhibition impinging on several distal basal dendrites could finely modulate the respective branch-specific NMDA spikes and, consequently, the number of Na^+^ spikes in the axon (Figure 7). This finding is consistent with the recent *in*-*vitro* and *in*-*vivo* findings reported in (Royer et al. 2012; Lovett-Barron et al. 2012).

### Dendritic inhibition controlling NMDA-induced plasticity in clusters of adjacent dendritic spines

NMDA spikes are known to be involved in facilitating long term potentiation (LTP) and depression (LTD) (Gambino et al. 2014; Sandler et al. 2016; Gordon et al. 2006; Brandalise et al. 2016; Golding et al. 2002). We showed in Figure 6 that precisely timed inhibition could gradually reduce the NMDA-dependent current influx by up to 60% of its original charge simultaneously in a group of spines cooperatively involved in generating the NMDA spike. Considering the Ca^2+^-control theory for inducing plasticity (Shouval et al. 2002; Graupner & Brunel 2012) and see also (Bar-Ilan et al. 2013), this implies that weak local dendritic (rather than spine) inhibition powerfully affects plasticity collectively in all spines involved in generating the NMDA spike. This capability to control plasticity in a specific group of spatially clustered dendritic spines by the relationship of inhibition and NMDA spikes adds to previous studies that have shown the impact of inhibition on plasticity induced by the back-propagating action potential (BPAP) and by dendritic Ca^2+^ spikes (Wilmes et al. 2016; Müllner et al. 2015; Pérez-Garci et al. 2013). It is important to note that the BPAP is relatively brief compared to the NMDA spine, so that inhibition should be more precisely timed if it interacts with BPAP. The Ca^2+^-spike, however, is long-lasting and comparable to the time-scale of the NMDA-spike, but the multiple voltage-dependence channels involved in generating the Ca^2+^-spike suggest that it cannot be finely tuned by inhibition in qualitatively the same way as the NMDA spike (Figure S4). Another important point to consider is that the Ca^2+^-spike is a global phenomenon, ignited near the main apical branch in L5 and L2/3 pyramidal cells (Larkum et al. 1999), whereas the NMDA-spike is highly branch-specific, and occurs at multiple dendritic loci (Major et al. 2008), thus enabling local inhibition to have the capability to control dendritic excitability at multiple dendritic sites of the pyramidal neurons. This inhibitory effect over a whole branch extends the case where only a single excitatory spine synapses is active (and no NMDA-spike is generated), making the effect of inhibition on this particular spine very specific and powerful, as was previously shown theoretically (Koch & Poggio 1983; Segev & Rall 1988) and experimentally (Chiu et al. 2013).

Another effect that timed dendritic inhibition might have on global dendritic plasticity is to suppress the briefer NMDA-spikes in terminal branches with relatively low input resistance, keeping intact the longer-lasting NMDA spikes in dendritic terminals with large input resistance (Figure 4). This contributes to the shift in the distribution of the NMDA-spikes duration from broad unimodal to bimodal distribution due to sparse dendritic inhibition (Figure 5). Consequently, timed dendritic inhibition could act as a cellular mechanism protecting the neuron from the influence of shorter NMDA spikes that, perhaps, carry less information relevant to its overall computation. Thus, one may consider weak inhibitory input arriving at different dendritic branches as a filter, or as an additional threshold, allowing only branches with NMDA-spikes that are resilient enough to survive the weak timed inhibitory input to influence both the cell’s output and its plasticity processes.

### Fine-tuning of the neuron’s output by sparse dendritic inhibition interacting with distal NMDA spikes

The NMDA spike, even when occurring on a distal apical dendritic branch, significantly boosts the excitatory charge reaching the soma. Several such spikes occurring simultaneously at several dendritic branches could generate a burst of output spikes in the axon (Polsky et al. 2009; Milojkovic et al. 2004). This was indeed replicated in our theoretical study (Figure 7) where we also showed that the number of spikes in this burst could be finely modulated by weak spatially-distributed dendritic inhibition that targets specific dendritic branches that generate local NMDA-spike. This branch-specific dendritic inhibition could arise from different classes of inhibitory interneurons, each of which was shown to target different dendritic domains (Stokes et al. 2014; Markram et al. 2004; Fishell & Tamás 2014; Bloss et al. 2016; Ma et al. 2010; Klausberger 2009) and is activated at different times with respect to the excitatory input (Ma et al. 2010; Pouille 2001; Pouille et al. 2009; Wehr & Zador 2003) (Also note (Silberberg & Markram 2007) that describe how Martinotti cells target pyramidal cells during feedback inhibition in a delayed fashion). Recently, (Wilson et al. 2016) showed that in ferrets V1, the receptive fields of cortical neurons are strongly shaped by local nonlinear dendritic responses resulting from the activity of spatially clustered excitatory synaptic input. If the functional properties of neurons are indeed determined to a large extent by clustered synaptic activity invoking strong local dendritic nonlinearity (rather than by voltage fluctuations arising from sparse E/I balance), then branch-specific synaptic inhibition could very effectively modulate the functional / computational properties of the neuron (e.g., by shifting its receptive field). These results thus suggest a new angle to the growing number of studies highlighting the computational advantages of branch-specific dendritic nonlinearity (Milojkovic et al. 2004; Polsky et al. 2009; Larkum et al. 2009; Smith et al. 2013; Palmer et al. 2014; Lavzin et al. 2012; Poirazi & Mel 2001)

### Robustness of this work and other related studies

In this work we primarily used the augmented Jahr and Stevens model for the NMDA-current ((Jahr & Stevens 1990), and see Methods) as do many contemporary theoretical studies. However, recent experimental work (Vargas-Caballero 2004; Moradi et al. 2013; Kampa et al. 2004; Clarke & Johnson 2008) has led to a new and potentially more realistic model of the NMDA-current. We therefore confirmed our key results using alternative models for the NMDA spike (Figure S1) and found that the findings described in the present work were robust in a variety of NMDA models, namely that the bi-phasic vulnerability function remained. Do note that the two phases were less strongly separated in the original Jahr and Stevens model, as that model’s VI curve is more linear, which fits the mechanism we suggest at Figure 3. The magnitude of the effect of timed inhibition and its exact timing with respect to the NMDA plateau phase also changed depending on the specific model used.

In our research, we focused on in vitro environment with synchronized inputs, in order to be able to study the interaction between the NMDA, AMPA, GABA_A_ and leak currents without additional factors. However, true in vivo environments will have a much shunted membrane (Rapp et al. 1992) and synchronized inputs will likely arrive with some jitter. To check whether the bi phasic vulnerability function of the NMDA spike is robust under these conditions, we tested the case where the same inhibitory conductance arrived with some temporal jitter (Figure S7C-D), and the case where the cell membrane was shunted and its resting potential raised due to background activity (Figure S7A-B). The vulnerability was robust in both cases, although it was stronger at the in vivo case and weaker with additional jitter.

Notably, (Rhodes 2006) was the first to study the effect of timed-inhibition on the NMDA spike. In his study, the NMDA / AMPA conductance ratio was much larger than in our study (10/1 and 1/1, respectively) and the inhibitory conductance in his work was larger (2 nS instead of 1 nS). Consequently, the Rhodes study demonstrated the termination phase, but did not show the two-phase effect of inhibition on the NMDA-spike (Figures 1 and 3), nor did it explore the gradual effect of inhibition over the NMDA-related current influx or its effect on the output spikes.

In recent years several studies ((Iacaruso et al. 2017; Wilson et al. 2016)) have demonstrated that local segregation of co-tuned synapses on specific dendritic branches, combined with dendritic nonlinearities, play a key role in shaping the orientation selectivity of these neurons. Our work demonstrated that dendritic inhibition targeting local dendritic branches could effectively control branch-specific nonlinearity; it is thus expected to strongly influence the global computation (e.g., the receptive field) of the neuron. This prediction is yet to be examined experimentally. In addition, in the hippocampus, work on neuronal oscillations has led to the hypothesis that temporal windows for computation and integration of inputs are determined by oscillations of interneuron activity (see (Mizuseki et al. 2009)). Our work may shed light on how temporally precise inhibition may bound such windows of input integration.

Overall, this study shows that during its long-lasting plateau phase and as a function of its unique dynamics, the dendritic NMDA spike is highly sensitive to well-timed synaptic inhibition. A relatively weak GABAergic inhibition can potently control the voltage dynamics and Ca^2+^ current influx generated by local dendritic NMDA spikes at their multi-site, mostly distal dendritic loci of origin. Consequently, distal dendritic inhibition could finely tune both branch-specific dendritic plasticity as well as the output spiking of the neuron. The NMDA spike combined with local dendritic inhibition may provide an adjustable “knob” for controlling the I/O repertoire of neurons and adjusting local dendritic plasticity at distal dendritic branches.

#### Future work

This theory was tested in a simulation in a biophysical model, and was not tested by us experimentally. In order to fully explore these results in vitro, we would suggest a combination of glutamate uncaging on the dendrites to account for the excitation, and either GABA_A_ uncaging or voltage clamping the soma to simulate the inhibition. It can be seen in Figure S2C that voltage clamping the dendritic membrane to a hyperpolarized state can reproduce the distinct recovery and termination of the NMDA spike after timed inhibition, and as can be seen in Figure S2A-B, this effect can be seen when recording from the soma, and should be visible with the existence active channels as well (Figure S2D).

In this work, we focused on the effect of timed inhibition on NMDA spikes, with special attention given to the local excitatory and inhibitory currents with respect to the local passive current and input resistance, either in the dendritic shaft or in the spine head. We did not venture into the effect of passive axial currents flowing to and from other dendrites, nor into the effect of other active channels such as voltage gated potassium channels or voltage gated calcium channels. These of course have an additional effect, which should be examined further, both in theoretical work and experiments. Specifically, we are interested how this time-dependent vulnerability of the NMDA spike changes with respect to other non-linear phenomena (specifically the Ca^2+^ spike, as can be seen in Figure S4), and how it is modified by varying conditions of active ion channel conductances (shown in Figure S2D, but not yet rigorously studied), for example the outward potassium current which might lend support to the hyperpolarization.

## Methods

### NEURON Simulation

The models and simulations were written and carried out in the NEURON simulator, wrapped with python script. All the models can be found in ModelDB, with scripts to recreate the figures. We chose to use several models in order to explore and characterize the effect of timed inhibition on the NMDA spike, for several reasons: The full biophysical model (Hay et al. 2011) which contains the passive, active, and morphological parameters that may affect the vulnerability of the NMDA spike is used to show that the effect is qualitatively present in this model (Figures 5, 7, S2D and S7A-B). However, in order to systematically study the individual components of the model, we chose to study the effect of the placement of the synapses without active properties in a passive model (Figures 1, 2, 4, S1, S2A-C and S7C-D). In addition, in order to fully characterize the dynamic between the excitatory NMDA current, the inhibitory GABA_A_ current and the passive leak current, we chose to study the intersections of the currents and the dynamics of the threshold in an isopotential model (Figures 3, S3, S4 and S5). For the spine head inhibition in Figure 6, we used the CA1 pyramidal cell model of (Grunditz et al. 2008).

### Morphological model

For a morphological model of the neuron, we used the Hay model of a layer 5 pyramidal cell of an adult rat (Hay et al. 2011). For Figures 1, 2, 4, S1, S2A-C and S7C-D we removed the active voltage dependent channels from the model, such that the model was passive aside from the NMDA synapses. For Figures 5, 7, S2D and S7A-B we used the full nonlinear model with all active channels. Excitatory synapses were placed on a distal apical branch (Figures 1, 2, 4 and 5) or basal terminal branch (Figures 4, 5 and 6), distributed with 1 excitatory synapse per 2.2 μm on the middle 44 μms of the branch (for Figures 1 and 2) or 1 μm on the last 20 μms of the branch (for Figures 4, 5 and 6). Inhibitory synapses were placed in the center of the distribution of the excitatory synapses, except in Figure 2, where their location was either in the center, 35 μms distal to the center, or 35 μms away from the center of the distribution of excitatory synapses.

### Single compartment model

To test in a more controlled environment without the effects of morphology and dendritic load, we used a single compartment model of an isopotential cell containing only the leak channels and the synapses for Figures 3, S3, S4 and S5. The model consisted of a soma 20 μm wide and 20 μm long, with a Rm of 20000, and a resting potential of -80 mV.

### Synaptic models

We used the Jahr and Stevens model (Jahr & Stevens 1990) for the NMDA current, with slight modifications. The gamma parameter, which defines the non-linearity of the magnesium dependence, was raised from 0.062 to 0.08, as was done in other studies (Rhodes 2006; Poleg-Polsky 2015; Lavzin et al. 2012). The bi-phasic response of the NMDA spike to inhibition was also demonstrated for other recent models of the NMDA current (Figure S3). The GABA_A_ and AMPA currents were modelled as double exponents, with the GABA_A_ rise and decay times at 0.18 and 5 milliseconds respectively (Gupta et al. 2000; Salin & Prince 1996), the AMPA rise and decay time set to 0.2 and 1.7 milliseconds respectively (Spruston et al. 1995), and the NMDA rise and decay time at 2 and 75 milliseconds respectively (Rhodes 2006; Jahr & Stevens 1990). The GABA_A_ reversal potential was -80, and the NMDA and AMPA reversal potential was 0. The NMDA / AMPA ratio was 1/1, as was used in (Hay et al. 2011).

The equation used for the simulation of the NMDA current in the modified Jahr and Stevens model was:

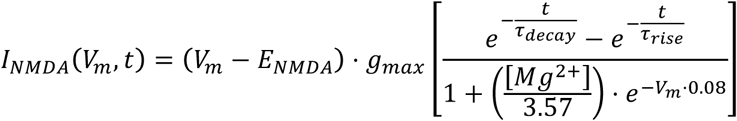

### Excitatory and Inhibitory interaction

To test the effect of inhibition on the NMDA spike when arriving at different times, we activated the excitatory synapses simultaneously, crossing the threshold and generating a dendritic NMDA spike. The GABA_A_ synapses were activated at different times, across a parameter space of 15 milliseconds before excitation and 80 milliseconds after excitation, thus sampling the parameter space every one millisecond in the vulnerability function graphs (Found in Figures 1, 2, 7, S1, S2, S3, S4, S6 and S7). Voltage was recorded from the center of the excitatory synapse distribution on the branch in the morphological model, and from the middle of the compartment in the isopotential cell model. The NMDA current was recorded from the sum of all excitatory synapses, the GABA_A_ current was recorded from the inhibitory synapse, and the leak current was recorded from the isopotential cell. For the calculation of the normalized voltage integrals, we summed the voltage after adding the resting potential to have a positive signed result, and divided by the voltage integral of the control case. For the current integral we have done the same, without adding the minimum value, and while taking the absolute value of the current. The summation covered the period from 10 milliseconds before the excitation to 200 milliseconds after the excitation, where the voltage of the NMDA spike almost vanished entirely.

### V-I curve

To create the VI graphs shown in Figure 3, we took the following approach: For each time point, we simulated the activation of the synapses up to that time point, then activated the voltage clamp to each voltage between -90 and 0, with a sampling of 0.1 millivolts, and recorded the currents 1 millisecond after the clamp started. A separate simulation was run for each voltage. The outward currents were multiplied by -1 for easier comparison with the inward currents. Because the time constant of the voltage change was faster than the time constant of the change in NMDA current, we considered the voltage change as instantaneous for the fixed point calculation. We defined the intersections between the V-I curve of the excitatory current and the negative of the outward current as fixed points, since when the voltage is at that location, it does not change, and if it is perturbed outside the fixed point it either diverges away from it or converges towards it. To create the trajectory of the fixed points over time (Figure 3B, S5), we recorded the intersections of the NMDA and leak currents after an activation of 8 nS of NMDA conductance, and plotted them over time. The voltage trace in these figures was created using the same excitatory and inhibitory conductances and activation times, without using the voltage clamp.

### Simultaneous activation of several dendritic branches

In Figure 7, 20 AMPA and NMDA synapses (0.4 nS each) were evoked on 16 different basal dendritic terminal branches uniformly distributed on the last 20 μm of each dendrite. Two inhibitory synapses (1 nS each) were placed in the center of the distribution on each branch. For Figure 7D, 60 possible combinations of 16 unique different basal dendritic terminal branches were randomly selected, such that each trial recorded the activation of one combination of 16 branches. The voltage was recorded from the soma for the spike statistics.

### NMDA spike initiation conductance threshold calculation

To calculate the amount of NMDA and AMPA conductance required to initiate the NMDA spike (for Figures S5), we activated increasing amounts of excitatory conductances, measured the voltage integral of the NMDA spike in each condition, and calculated the derivative of the integral / conductance function. The excitatory conductance creating the maximum of this derivative function was considered to be the initiation conductance threshold of the NMDA spike.

### NMDA spike termination conductance threshold calculation

To calculate the amount of GABA_A_ conductance required to terminate the NMDA spike (for Figures 4 and S5), we activated increasing amounts of inhibitory conductances, measured the voltage integral of the NMDA spike in each condition, and calculated the derivative of the integral / conductance function. The inhibitory conductance creating the minimum of this derivative function was considered to be the termination conductance threshold of the NMDA spike. When no threshold was calculated for the termination (Figures 1-3 and 5-7) premature termination was defined as the NMDA spike half-width being less than 90% of the control case.

### Histogram of NMDA spike lengths

In Figure 5, we activated 20 excitatory synapses on each terminal dendrite (similar to Figure 5) in a full nonlinear model recording the voltage from the center of the distribution on the branch. To account for varying excitatory strengths, we sampled the NMDA conductance between 0.1 nS and 1 nS at steps of 0.1 nS. The histograms in Figure 5 show all the NMDA spikes evoked using this stimulation on all terminal dendrites, with all the tested conductances. An NMDA spike was defined as an event where the local dendritic voltage was above -40 mV for at least 20 msecs following excitation, in the control case without inhibition.

### [Ca^2+^] concentration and voltage recording on spine head

In figure 6, we used the CA1 pyramidal cell model of (Grunditz et al. 2008) to obtain an accurate description of the voltage and [Ca^2+^] concentration on the spine head. We remove the R-type calcium channels from the model to concentrate on the effect of spine morphology on the interaction between timed inhibition and excitation. The kinetics of Ca2+ influx and diffusion, as well as the NMDA and AMPA models in this Figure were taken from (Grunditz et al. 2008).

## Supplemental material

**Supplemental Figure 1.**
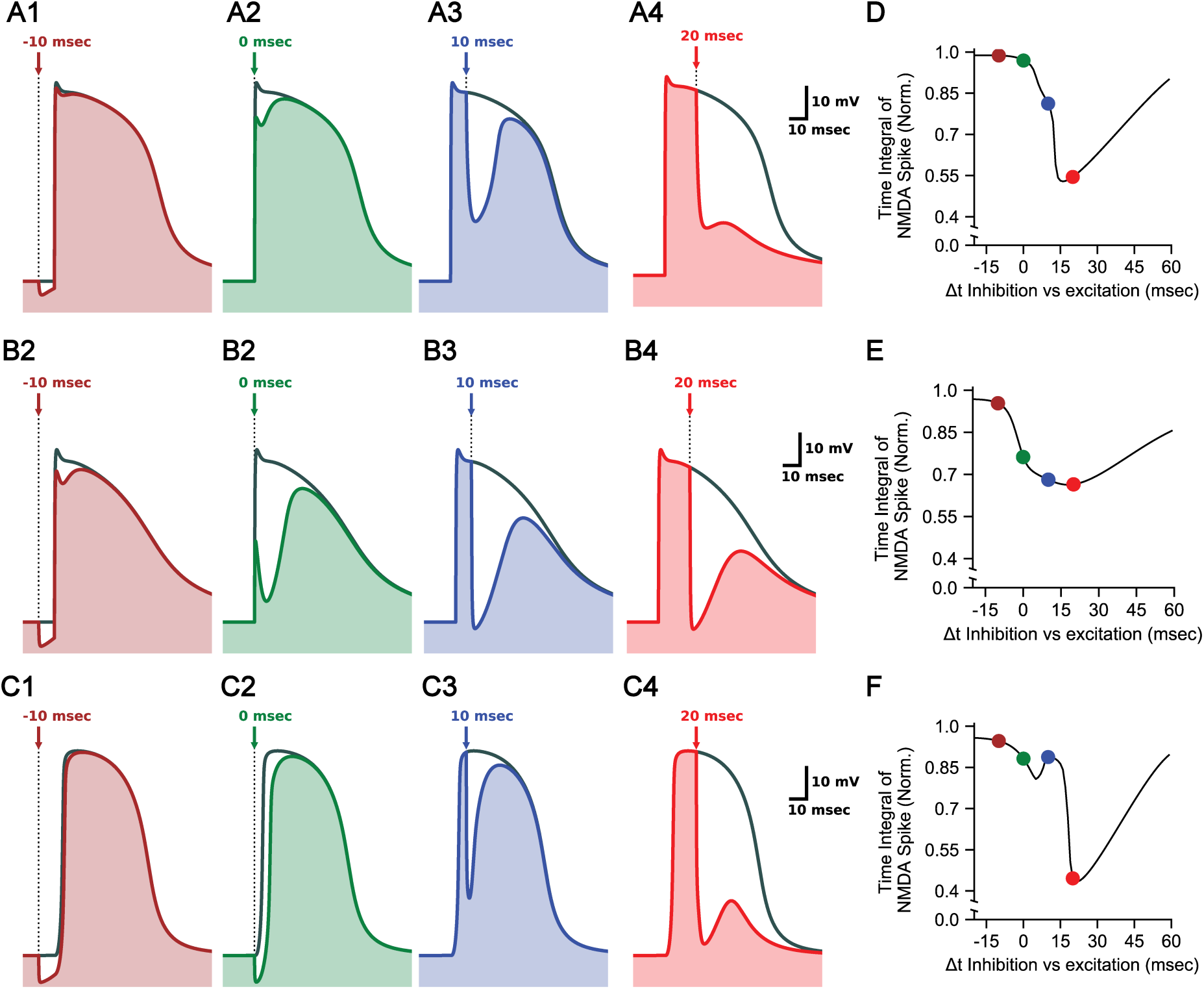
The phenomenon is robust in other NMDA receptor models. (A)-(C). Similar to Figure 1B, using different NMDA receptor models (see Supplemental Methods for model descriptions), and with other ratios of excitation-inhibition. (A1-4). Similar to Figure 1B, using the modulated Jahr and Stevens model, with 20 AMPA and NMDA synapses of 0.4 nS each, and a single GABA_A_ synapse with 1 nS. Inhibition arrived at -10, 0, 10 and 20 msec, as indicated by the arrows in each figure. (B1-4). Similar to A, with the unmodulated Jahr and Stevens model. In this case, more inhibition and less excitation were required to reproduce the phenomenon (12 excitatory and 12 inhibitory synapses). (C1-4). Similar to A, with the Vargas-Caballero and Robinson model. In this case, slightly less inhibition was required (0.7 GABA_A_ nS instead of 1 GABA_A_ nS).

**Supplemental Figure 2.**
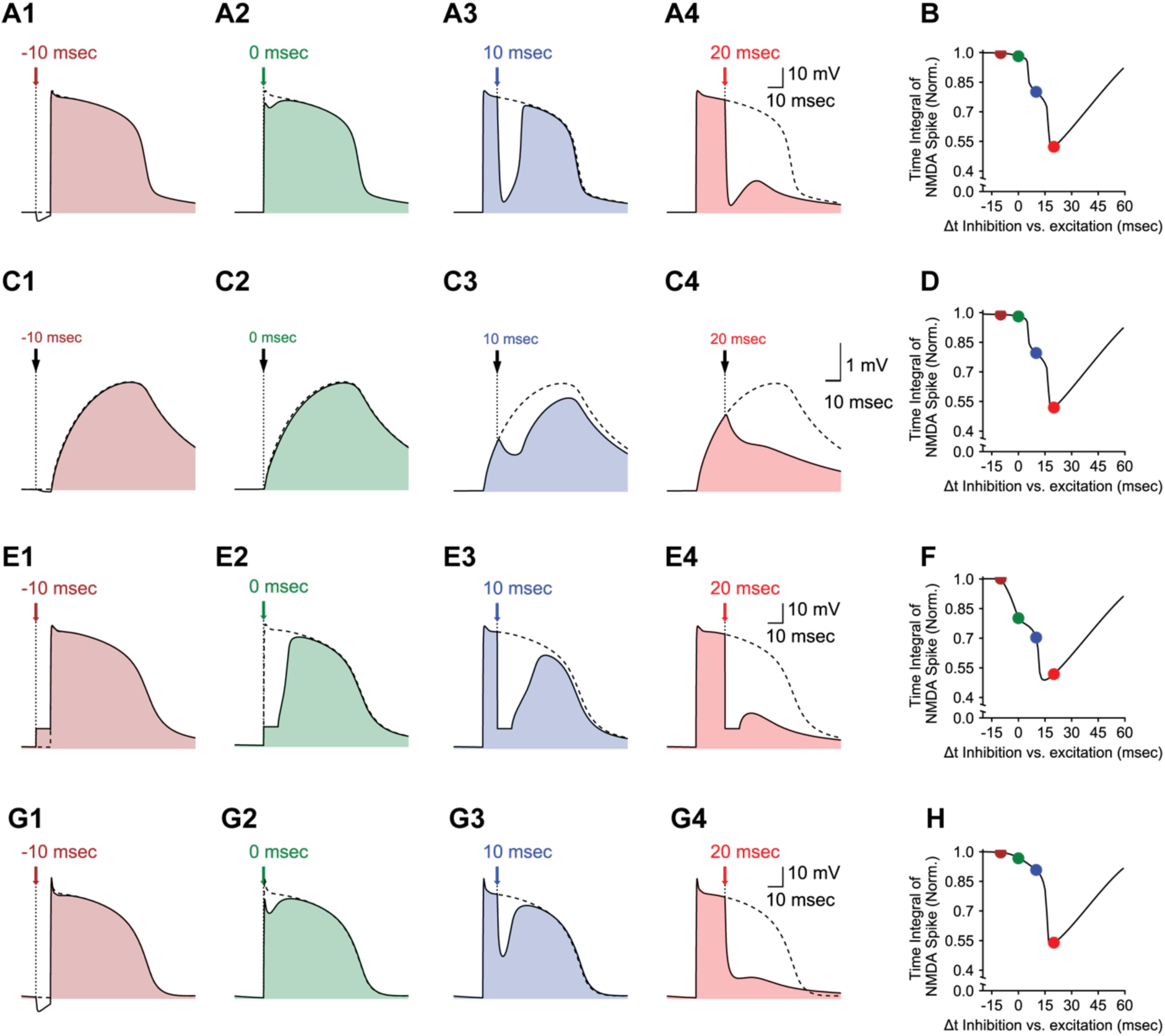
The phenomenon is robust across different neuron models. Similar to Figure 1B, with different experimental frameworks and neuron models. (A1-4) On a reconstructed L5 pyramidal neuron biophysical model without active channels, 22 excitatory synapses and one inhibitory synapse located on a proximal basal dendrite 186 μm from the soma. Voltage recorded from dendrite. (B) As in Figure 1D, the vulnerability function of the case in (A). (C1-4) As in (A), with voltage recorded from soma. (D) The vulnerability function of (C). Shape is near identical to that of (B) (E1-4) As in (A), without GABA_A_ synapses, and with voltage clamped to -60 mV from the onset of the inhibition (colored arrow) to 10 msec afterwards. (F) The vulnerability function of (E). Shape is similar to that of (B) and (D) (G1-4) On a reconstructed L5 pyramidal neuron biophysical model with active channels, and with 19 instead of 20 AMPA- and NMDA-based excitatory synapses. (H) The vulnerability function of (G). Shape is similar to that of (B), (D) and (F).

## Early weak inhibition increases, rather than decreases, the influx of NMDA current

Analysis of the V-I curve of the NMDA-mediated dynamics indicated that the peak (the maximal deep) of the NMDA current was at a more hyperpolarized value than that of the “upper stable” fixed point (e.g., see Figure 3B red trace). This means that a certain magnitude of inhibition, although hyperpolarizing the NMDA plateau, would actually increase the NMDA-mediated current. This case is shown in Figure S3A where the NMDA spike was transiently hyperpolarized by some 15 mV (Figure S3A, top) but, due to an increase in the driving force, the NMDA current increased more than the control levels (Figure S3A, dark red at bottom). To study the influence of timed inhibition on the NMDA current, we activated the inhibition at different times along the course of the NMDA spike while recording both the NMDA gated voltage and current. As expected from the argument above, early weak inhibition hyperpolarized the NMDA gated voltage such that the resultant NMDA current was larger than in the control case (without inhibition, Figure S3A). Late weak inhibition (Figure S3B), or stronger early inhibition (not shown) hyperpolarized the voltage to a region where the voltage at the NMDA related V-I curve was lower than in the control case and thus no extra NMDA current was generated due to inhibition. The maximal amount of NMDA current that could be reached due to inhibition riding on the plateau phase of the NMDA spike is shown, by the blue vertical line in Figure S3C.

To further quantify this effect, we calculated the integral of the current for different inhibition arrival times and normalized it by the integral of the current without inhibition. Figure S3D shows that the current integral was larger than the control level for inhibition activated at early times (shaded red area). At later times the NMDA related current decreased because the voltage reached regions where the NMDA current was lower. In contrast, the voltage time integral was smaller than the control throughout the inhibition arrival time range examined here (Figure S3D, blue curve). For the range of parameters used here, the integral of the NMDA current could grow by up to 13% as compared to the control case. Finally, we explored the range of this paradoxical boosting effect by inhibition for a range of parameters, and created a heat map as shown in Figure S3E. These findings suggest that during the plateau phase of the NMDA spike, weak dendritic inhibition could increase, rather than decrease the flow of inward current (including Ca^2+^ current) into the dendrites, possibly enhancing synaptic plasticity in this dendritic branch (see Discussion).

**Supplemental Figure 3.**
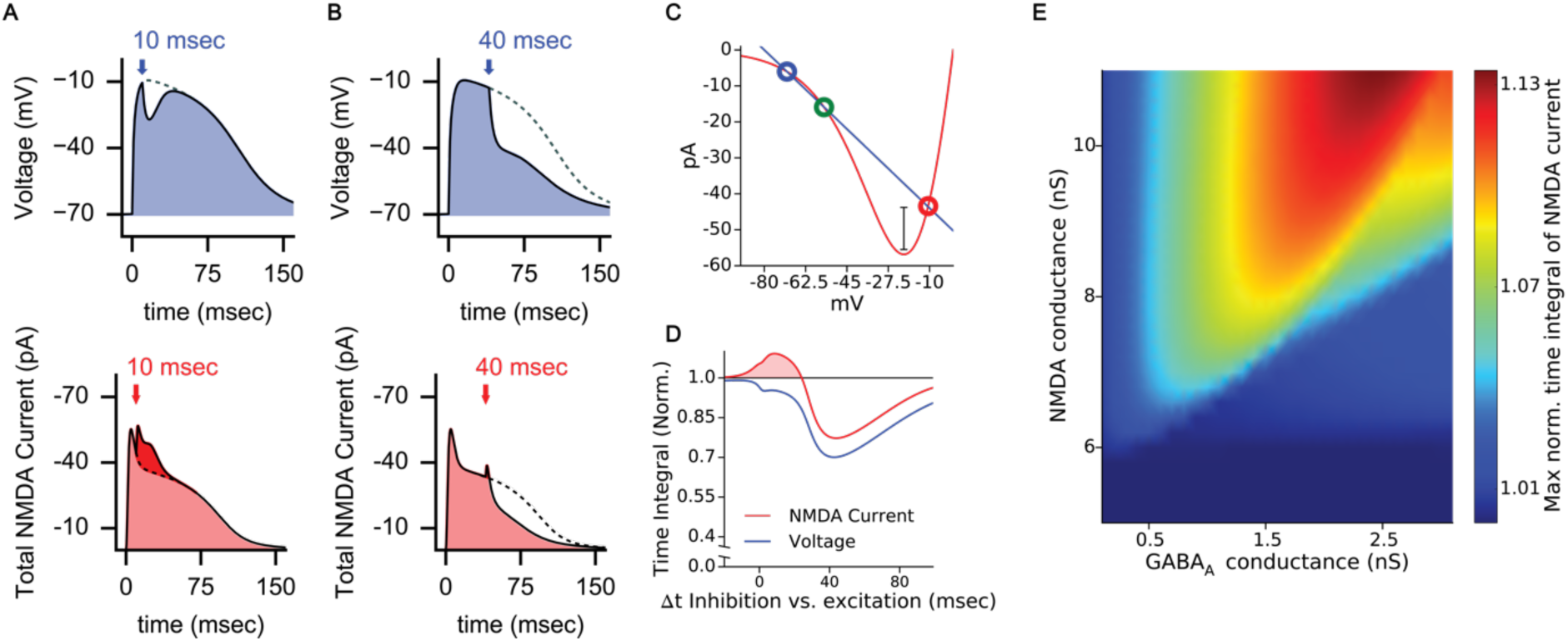
Early inhibition may increase, rather than decrease, the influx of NMDA current. (A) Top. As in Figure 1B with inhibition arriving at Δt = 10 msec. Bottom: The NMDA current under the same conditions as at the top. The dashed line depicts the NMDA current in the control case. The continuous line (and the corresponding light red shaded area) show the current trace with inhibition; the additional inward current due to inhibition is denoted by the dark red shaded area. (B) As in (A), but with inhibition arriving at Δt = 40 msec. (C) The V-I curve of the NMDA spike, 5 milliseconds after excitation onset. Blue and red curves and colored circles are as in Figure 3A. The vertical line above the minimum of the red curve shows the maximal additional NMDA current due to early weak inhibition. The length of this line is the difference between the NMDA current at its deepest point and that obtained at the right-most fixed point (red circle). (D) Same as in Figure 1D, for both the normalized voltage time integral (blue) and the normalized current integral (red). Early weak inhibition increases the total influx of NMDA current above control levels (shaded red area). In (A)-(D) the model used was as in Figure 1A. (E) The maximum increase in NMDA current due to early inhibition for different GABA_A_ and NMDA conductances in a model of an isopotential neuron. In this case, early inhibition increased the NMDA current by about 13% (dark red region).

**Supplemental Figure 4.**
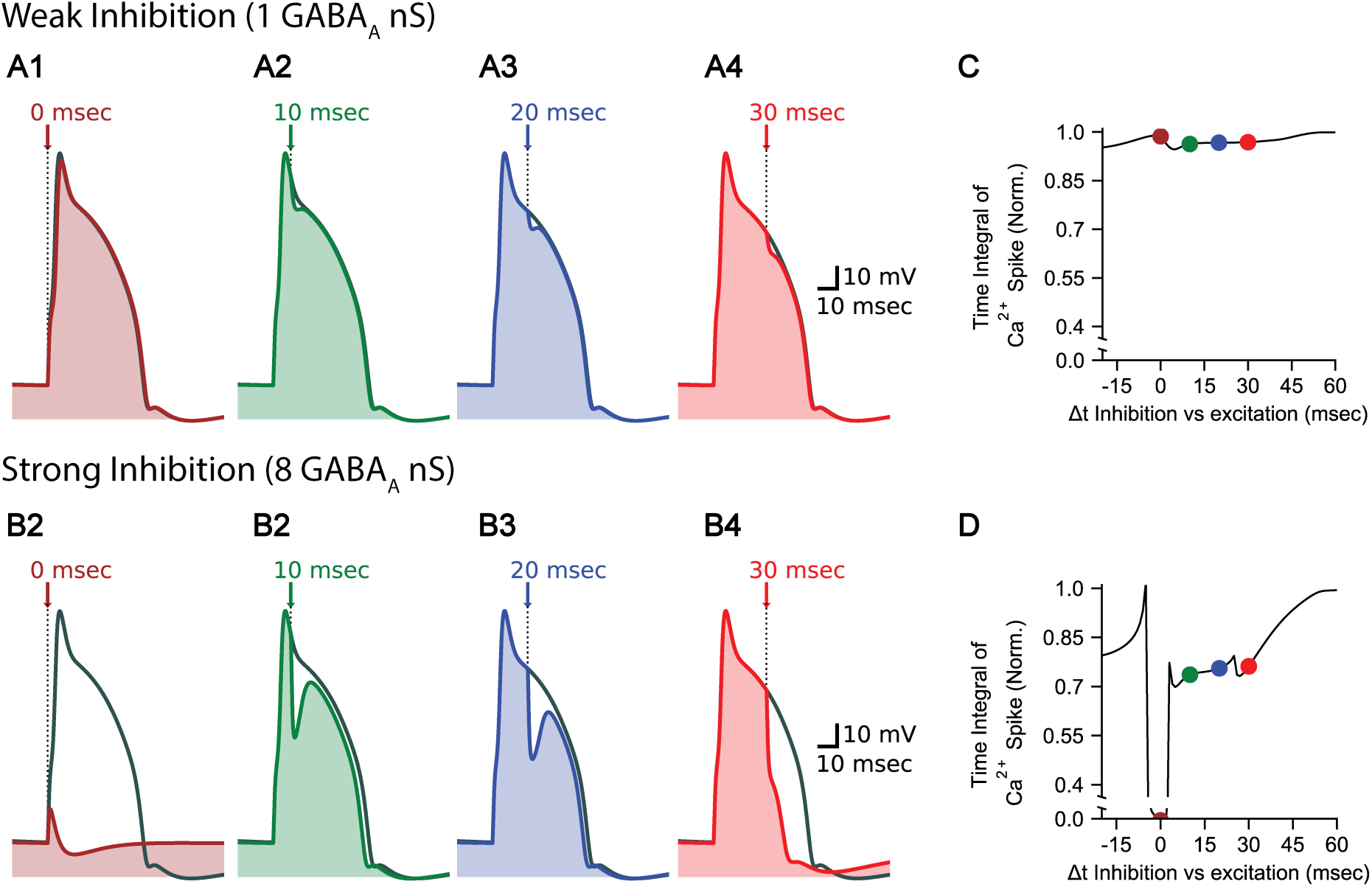
Ca spikes exhibit qualitatively different vulnerability to timed inhibition, and are generally more resilient. (A) Weak inhibition does not affect Ca^2+^ spike in the same way as it affects NMDA spikes. 20 AMPA synapses (without NMDA component) were activated simultaneously in an isopotential cell with active properties similar to those in the nexus of the L5 pyramidal cell (see Supplemental Methods). A single GABA_A_ synapse with a conductance of 1 nS was activated at different times (similar to Figure 1B). (B) Similar to Figure S4A, with 8 GABA_A_ nS. Strong inhibition terminated the Ca spike when arriving at the excitation onset, but did not terminate it prematurely when arriving 10 and 20 milliseconds after excitation onset. Strong late inhibition arriving 30 milliseconds after excitation onset terminated the Ca spike prematurely. (C) Weak inhibition hardly influences the Ca^2+^ spike. Similar to Figure 1C, the vulnerability function of the Ca^2+^ spike with GABA_A_ conductance of 1 nS. The black curve is the integral of the Ca^2+^ spike with inhibition arriving at different times, divided by the integral of the Ca^2+^ spike without inhibition. The colored dots correspond to the traces on Figure S4A. (D) Strong inhibition effects Ca^2+^ spikes differently than NMDA spikes. Similar to Figure S4C, with GABA_A_ conductance of 8 nS. The colored dots correspond to the traces on Figure S4B.

**Supplemental Figure 5.**
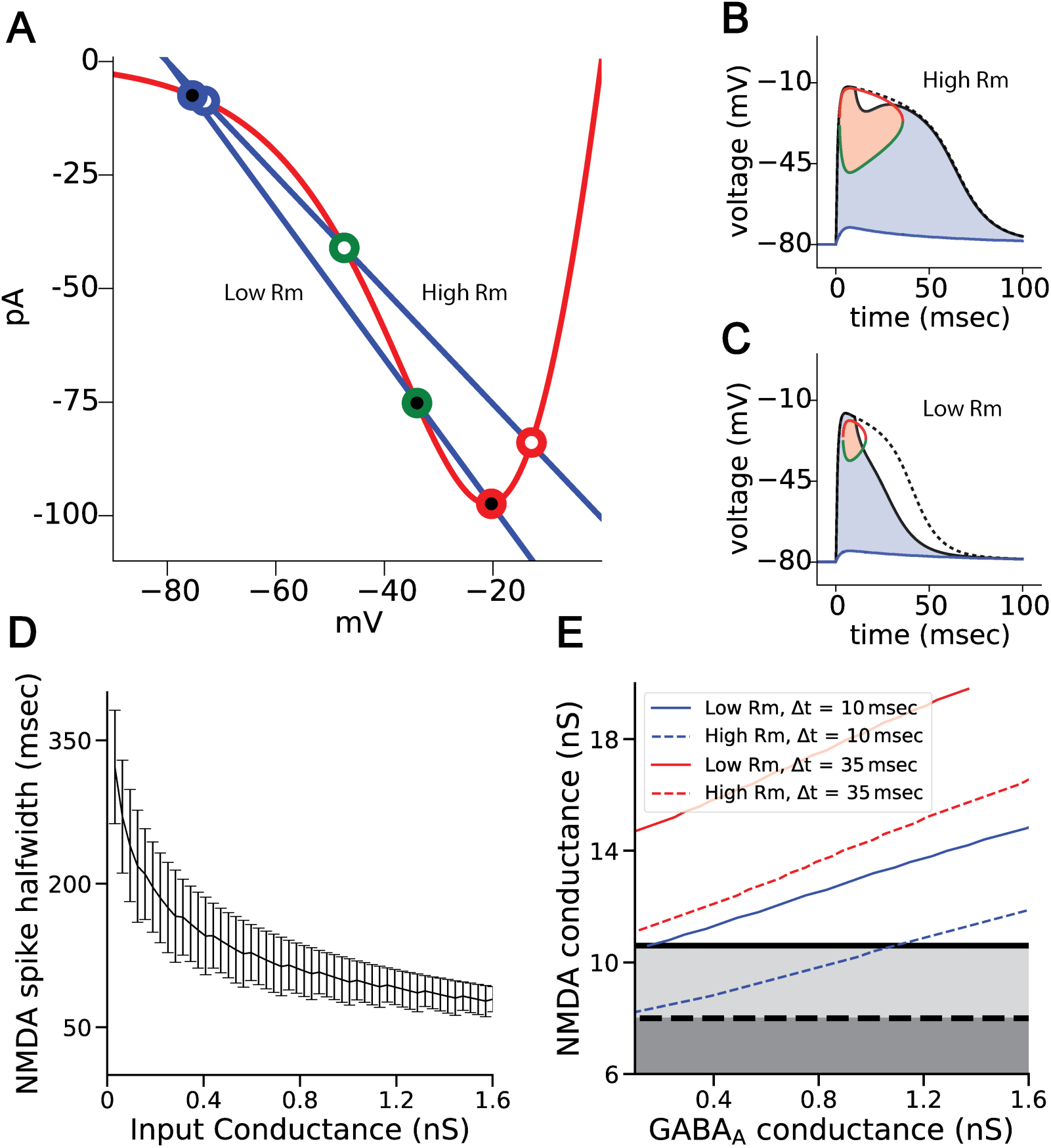
Higher input resistance decreases the initiation threshold, increases spike height and length, and makes NMDA spikes more resilient to inhibition. Model as in Figure 3 in the main text. (A) Similar to Figure 3A, with the red curve being the NMDA current, the two blue curves being the larger and smaller leak currents, corresponding to lower and higher input resistances. Lower input resistance acts as additional outward current, depolarizing the initiation threshold (blue open dots) thus making the initiation harder, and hyperpolarizing the upper fixed point (green open dots), and effectively decreasing the maximal NMDA spike height and length. White filled dots have lower input resistance; black filled dots have higher input resistance. (B) Similar to Figure 3B. Higher input resistance allows for more resilient NMDA spikes. Inhibitory input of 1 nS *GABA_A_*. conductance at 10 milliseconds drops the voltage to above the threshold, allowing the NMDA spike to recover after the inhibition. (C) Similar to Figure 3B. Lower input resistance makes for lower and shorter NMDA spikes. Inhibitory input of 1 nS *GABA_A_*. conductance at 10 milliseconds drops the voltage below the threshold, causing the NMDA spike to terminate after the inhibition. (D) The NMDA half-width rises with input resistance (and decreases with input conductance). The black curve is the mean NMDA half-width, with error bars measured over different NMDA conductances. (E) Response function of the NMDA spike to timed inhibition changes with input resistance. Dashed lines represent high input resistance, solid lines represent low input resistance. Black lines represent the threshold of NMDA spike below which an NMDA spike cannot be evoked. Blue lines represent the amount of *GABA_A_* conductance needed to terminate an NMDA spike created with a specific conductance with an delay of 10 milliseconds between excitation and inhibition. Red lines are identical but with delay of 40 millisecond.

**Supplemental Figure 6.**
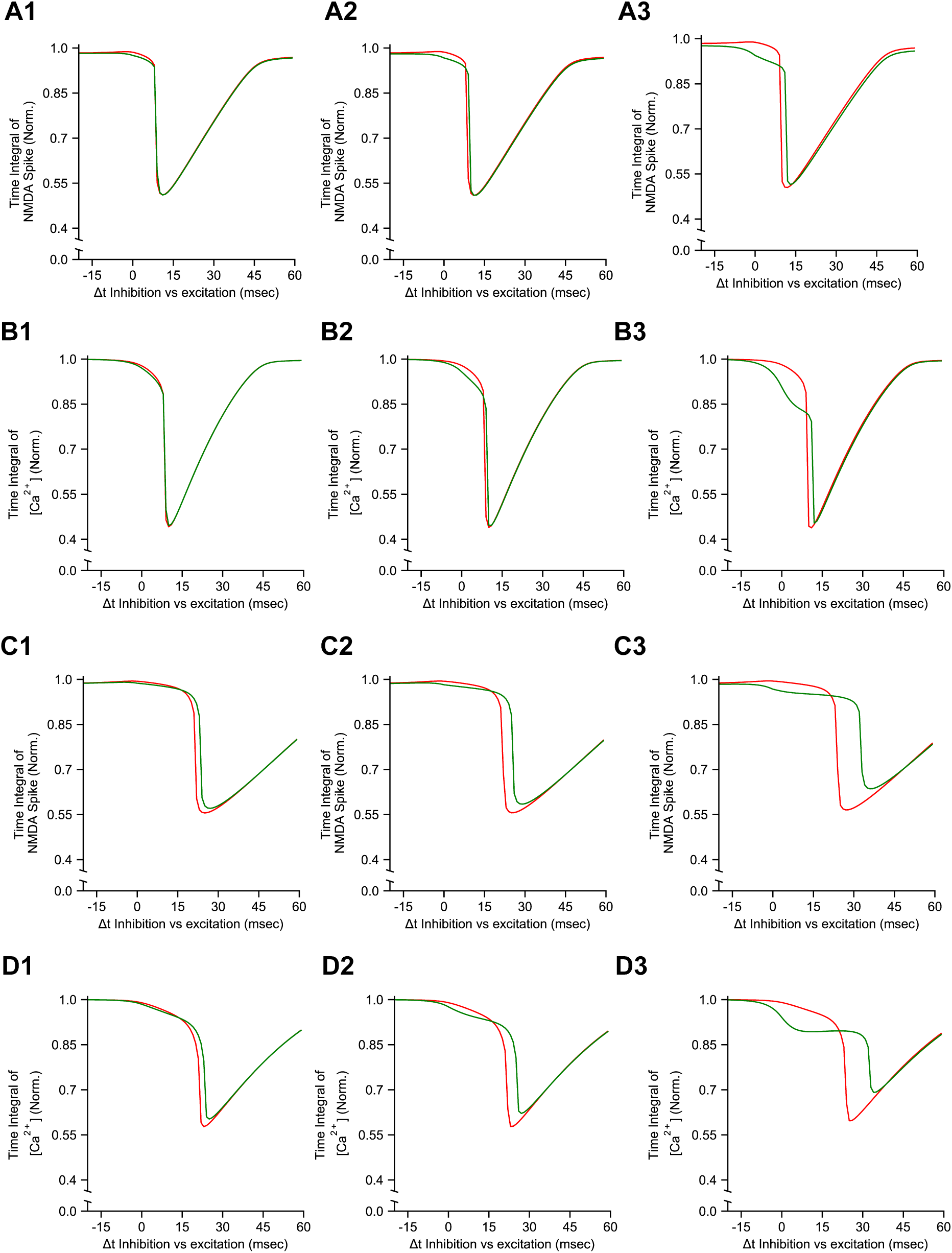
Larger spine neck resistance and larger branch membrane resistance decreases the effect of spine inhibition. Model as in Figure 6 in the main text. (A1-3) Similar to Figure 6G. Vulnerability function of the NMDA spike voltage recorded from a single spine when inhibition is located at the spine head (green) or at the dendrite (red). Excitation and inhibition were placed on a proximal basal dendrite with input resistance of 1583 *MΩ*. (A1) is the case where spine neck resistance was 120 M*Ω*. (A2) is the case where spine neck resistance was 600 M*Ω*. (A3) is the case where spine neck resistance was 1.2 G*Ω*. (B1-3) Similar to (Figure 6H). Vulnerability function of the total calcium concentration [Ca^2+^] in a single spine when inhibition is located at the spine head (green) or at the dendrite (red). Excitation and inhibition were placed on a proximal basal dendrite. Spine neck resistances in (B1-3) correspond to those in (A1-3). (C1-3) Similar to (A), with excitation and inhibition placed on a distal apical dendrite with input resistance of 1956 *MΩ*. Spine neck resistances in (C1-3) correspond to those in (A1-3). (D1-3) Similar to (B), with excitation and inhibition placed on a distal apical. Spine neck resistances in (D1-3) correspond to those in (A1-3).

**Supplemental Figure 7.**
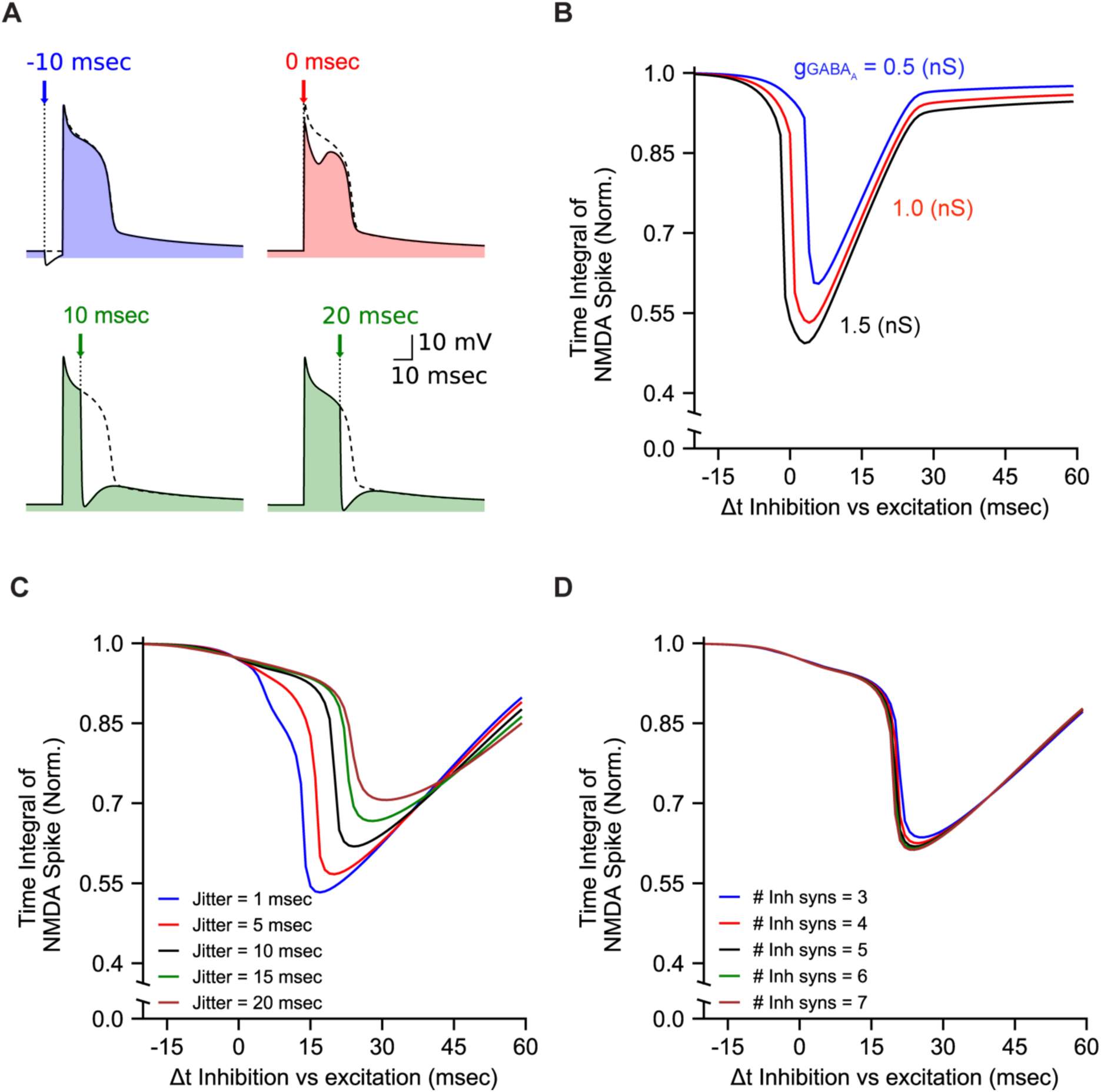
The two phasic vulnerability decreases yet stays robust when jitter is added to the inhibitory synapses, and increases under a simulated in vivo background activity shunt. (A) Similar to Figure 1B, with a higher leak conductance and leak current reversal potential simulating in vivo background activity (20 excitatory synapses with 0.4 nS conductance each, distributed around the last 20 μm on a distal apical dendrite, 1 inhibitory GABA_A_ synapse in the middle of the distribution). (B) Similar to Figure 1D, with the same conditions as in A. (C) Wider temporal distribution decreases the vulnerability function. Similar to Figure 1D, with 5 weak (0.2 nS) inhibitory synapses arriving in uniform temporal distribution around *Δt*. (D) Distribution of the inhibition conductance across more synapses does not change the vulnerability function. Similar to Figure 1D, with a number of inhibitory synapses (whose sum of conductance is 1 nS) arriving in a uniform temporal distribution around 10 Δt.

## Supplemental methods

The full biophysical model used in figure S1 is the (Hay et al. 2011) model of the L5 rat pyramidal cell, including the active channels.

The augmented Jahr and Stevens model used in this paper and in figure S1A is based on the Jahr and Stevens experiments (Jahr & Stevens 1990), with the gamma parameter changed from 0.062 to 0.08, in order to create the nonlinearity of the NMDA spike. Although the change from 0.062 to 0.08 is arbitrary, it is the model used in most recent modeling papers. To account for its arbitrariness, we used other models to check its robustness in more experiment based models. The original Jahr and Stevens model used in figure S1B had a time dependent component and a voltage dependent component to account for the time kinetics of the receptor from the onset of glutamate binding, the voltage dependency of the magnesium block, and the driving force. In this model, the gamma parameter of the magnesium block remained at 0.062 as was shown in the 1990 experiment, which allows for less nonlinearity, which compromises the threshold nature of the NMDA spike. The Vargas-Caballero and Robinson model was created in 2005 based on the work of Vargas-Caballero (Vargas-Caballero 2004) who systematically characterized the properties of the NMDA spike kinetics. This 10 state kinetic model accounts for more detailed properties of the NMDA receptor current, such as the temperature dependency of the time constants, non-magnesium related voltage dependency, and desensitization of the NMDA receptors to reoccurring input. Despite the differences, nonlinearity persists in this model as well, and our results are qualitatively very similar (compare Figures S1D and S1F).

The calcium spike in Figure S4 was created in an isopotential cell with a diameter and length of 20 μM, a Rm of 20000 MΩ, and active channels simulating the nexus of the layer 5 pyramidal cell in Hay’s model. The calcium spike was created using AMPA synapses to be able to compare to the NMDA spike created with the same AMPA synapses. No back propagation action potential was used, so that inhibition on the Ca spike could be tested without additional excitatory inputs such as bursting.

The in vivo background shunt simulation used in Figure S7 was designed as follows: The steady background conductance of 10000 excitatory and 2500 inhibitory synapses (0.4 nS and 1 nS each, respectively) was calculated and normalized for each branch, assuming a uniform distribution of synapses across the membrane length. The steady conductance was calculated using the methods of (Rapp et al. 1992), and added to the leak conductance, effectively producing the same shunt as in vivo background activity. The leak current reversal potential was set to be the mean voltage of activation of the 12500 synapses across the cell (-67 mV). The local excitatory and inhibitory synapses were placed on the last 20 μM of the dendrite.

## Acknowledgements

This work was supported by the Gatsby Charitable Foundation and the EPFL-Hebrew University Collaborative Grant, the EPFL support to the Laboratory of Neural Microcircuitry (LNMC), the ETH Domain for the Blue Brain Project (BBP), and by the Human Brain Project from the European Union Seventh Framework Program (FP7/2007-2013) under grant agreement no. 604102 (HBP).

## Author Contributions

M.D. and I.S. conceived the study and wrote the manuscript. M.D. carried out the analysis. G.C., E.M. and H.M. participated in discussions.

## Additional information

We declare no competing financial interests.

## References

Bar-Ilan, L., Gidon, A. & Segev, I., 2013. The role of dendritic inhibition in shaping the plasticity of excitatory synapses. Frontiers in Neural Circuits, 6(April), pp.1–13. Available at: http://journal.frontiersin.org/article/10.3389/fncir.2012.00118/abstract.

Bloss, E.B. et al., 2016. Structured Dendritic Inhibition Supports Branch-Selective Integration in CA1 Pyramidal Cells. Neuron, 89(5), pp.1016–1030. Available at: http://dx.doi.org/10.1016/j.neuron.2016.01.029.

Branco, T., Clark, B.A. & Häusser, M., 2011. Dendritic Discrimination of Temporal Input Sequences in Cortical Neurons. Science, 1671(2010), pp.1671–1676.

Branco, T. & Häusser, M., 2011. Synaptic Integration Gradients in Single Cortical Pyramidal Cell Dendrites. Neuron, 69(5), pp.885–892. Available at: http://linkinghub.elsevier.com/retrieve/pii/S0896627311001036.

Brandalise, F. et al., 2016. Dendritic NMDA spikes are necessary for timing-dependent associative LTP in CA3 pyramidal cells. Nature Communications, 7, p.13480. Available at: http://dx.doi.org/10.1038/ncomms13480.

Chiu, C.Q. et al., 2013. Compartmentalization of GABAergic Inhibition by Dendritic Spines. Science, 340(6133), pp.759–762. Available at: http://www.sciencemag.org/cgi/doi/10.1126/science.1234274.

Clarke, R.J. & Johnson, J.W., 2008. Voltage-dependent gating of NR1/2B NMDA receptors. The Journal of Physiology, 586(23), pp.5727–5741. Available at: http://doi.wiley.com/10.1113/jphysiol.2008.160622.

Cuntz, H., Remme, M.W.H. & Torben-Nielsen, B., 2014. The Computing Dendrite H. Cuntz, M. W. H. Remme, & B. Torben-Nielsen, eds., New York, NY: Springer New York. Available at: http://link.springer.com/10.1007/978-1-4614-8094-5 [Accessed May 7, 2017].

Fishell, G. – Tamás, G., 2014. Inhibition: synapses, neurons and circuits. Current Opinion in Neurobiology, 26, pp.v–vii. Available at: http://linkinghub.elsevier.com/retrieve/pii/S0959438814000713.

Gambino, F. et al., 2014. Sensory-evoked LTP driven by dendritic plateau potentials in vivo. Nature, 515(7525), pp.116–119. Available at: http://www.nature.com/doifinder/10.1038/nature13664.

Gidon, A. & Segev, I., 2012. Principles Governing the Operation of Synaptic Inhibition in Dendrites. Neuron, 75(2), pp.330–341. Available at: http://linkinghub.elsevier.com/retrieve/pii/S0896627312004813.

Golding, N.L. & Spruston, N., 1998. Dendritic Sodium Spikes Are Variable Triggers of Axonal Action Potentials in Hippocampal CA1 Pyramidal Neurons. Neuron, 21(5), pp.1189–1200. Available at: http://linkinghub.elsevier.com/retrieve/pii/S0896627300806352.

Golding, N.L., Staff, N.P. & Spruston, N., 2002. Dendritic spikes as a mechanism for cooperative long-term potentiation. Nature, 418(6895), pp.326–331. Available at: papers3://publication/doi/10.1038/nature00854.

Gordon, U., Polsky, A. & Schiller, J., 2006. Plasticity Compartments in Basal Dendrites of Neocortical Pyramidal Neurons. Journal of Neuroscience, 26(49), pp.12717–12726. Available at: http://www.jneurosci.org/cgi/doi/10.1523/JNEUROSCI.3502-06.2006.

Graupner, M. & Brunel, N., 2012. Calcium-based plasticity model explains sensitivity of synaptic changes to spike pattern, rate, and dendritic location. Proceedings of the National Academy of Sciences, 109(10), pp.3991–3996. Available at: http://www.pnas.org/cgi/doi/10.1073/pnas.1109359109.

Grunditz, A. et al., 2008. Spine Neck Plasticity Controls Postsynaptic Calcium Signals through Electrical Compartmentalization. Journal of Neuroscience, 28(50), pp.13457–13466. Available at: http://www.jneurosci.org/cgi/doi/10.1523/JNEUROSCI.2702-08.2008.

Gupta, A., Wang, Y. & Markram, H., 2000. Organizing principles for a diversity of GABAergic interneurons and synapses in the neocortex. Science (New York, N.Y.), 287(5451), pp.273–8. Available at: http://www.sciencemag.org/cgi/doi/10.1126/science.287.5451.273.

Hay, E. et al., 2011. Models of Neocortical Layer 5b Pyramidal Cells Capturing a Wide Range of Dendritic and Perisomatic Active Properties L. J. Graham, ed. PLoS Computational Biology, 7(7), p.e1002107. Available at: http://dx.plos.org/10.1371/journal.pcbi.1002107.

Hay, E. & Segev, I., 2015. Dendritic Excitability and Gain Control in Recurrent Cortical Microcircuits. Cerebral Cortex, 25(10), pp.3561–3571. Available at: https://academic.oup.com/cercor/article-lookup/doi/10.1093/cercor/bhu200.

Higley, M.J., 2014. Localized GABAergic inhibition of dendritic Ca2+ signalling. Nature Reviews Neuroscience, 15(9), pp.567–572. Available at: http://www.ncbi.nlm.nih.gov/pubmed/25116141.

Higley, M.J. & Sabatini, B.L., 2012. Calcium Signaling in Dendritic Spines. Cold Spring Harbor Perspectives in Biology, 4(4), pp.a005686–a005686. Available at: http://cshperspectives.cshlp.org/lookup/doi/10.1101/cshperspect.a005686.

Hoffman, D. a et al., 1997. K+ channel regulation of signal propagation in dendrites of hippocampal pyramidal neurons. Nature, 387(6636), pp.869–75. Available at: http://www.nature.com/doifinder/10.1038/43119.

Iacaruso, M.F., Gasler, I.T. & Hofer, S.B., 2017. Synaptic organization of visual space in primary visual cortex. Nature, 547(7664), pp.449–452. Available at: http://www.nature.com/doifinder/10.1038/nature23019.

Jack, J.J.B., Noble, D. & Tsien, R.W., 1975. Electric current flow in excitable cells, Clarendon Press. Available at: https://books.google.co.il/books?id=dsFqAAAAMAAJ.

Jadi, M. et al., 2012. Location-Dependent Effects of Inhibition on Local Spiking in Pyramidal Neuron Dendrites B. S. Gutkin, ed. PLoS Computational Biology, 8(6), p.e1002550. Available at: http://dx.plos.org/10.1371/journal.pcbi.1002550.

Jahr, C.E. & Stevens, C.F., 1990. Voltage dependence of NMDA-activated macroscopic conductances predicted by single-channel kinetics. The Journal of neuroscience, 10(9), pp.3178–3182.

Johnston, D. & Narayanan, R., 2008. Active dendrites: colorful wings of the mysterious butterflies. Trends in Neurosciences, 31(6), pp.309–316. Available at: http://linkinghub.elsevier.com/retrieve/pii/S0166223608001197.

Kampa, B.M. et al., 2004. Kinetics of Mg 2+ unblock of NMDA receptors: implications for spike-timing dependent synaptic plasticity. The Journal of Physiology, 556(2), pp.337–345. Available at: http://doi.wiley.com/10.1113/jphysiol.2003.058842.

Klausberger, T., 2009. GABAergic interneurons targeting dendrites of pyramidal cells in the CA1 area of the hippocampus. European Journal of Neuroscience, 30(6), pp.947–957. Available at: http://doi.wiley.com/10.1111/j.1460-9568.2009.06913.x.

Koch, C. & Poggio, T., 1983. A Theoretical Analysis of Electrical Properties of Spines. Proceedings of the Royal Society B: Biological Sciences, 218(1213), pp.455–477. Available at: http://rspb.royalsocietypublishing.org/cgi/doi/10.1098/rspb.1983.0051.

Larkum, M.E. et al., 2009. Synaptic Integration in Tuft Dendrites of Layer 5 Pyramidal Neurons: A New Unifying Principle. Science, 325(5941), pp.756–760. Available at: http://www.sciencemag.org/cgi/doi/10.1126/science.1171958.

Larkum, M.E. & Nevian, T., 2008. Synaptic clustering by dendritic signalling mechanisms. Current Opinion in Neurobiology, 18(3), pp.321–331. Available at: http://linkinghub.elsevier.com/retrieve/pii/S095943880800086X.

Larkum, M.E., Zhu, J.J. & Sakmann, B., 1999. A new cellular mechanism for coupling inputs arriving at different cortical layers. Nature, 398(March), pp.338–341.

Lavzin, M. et al., 2012. Nonlinear dendritic processing determines angular tuning of barrel cortex neurons in vivo. Nature, 490(7420), pp.397–401. Available at: http://www.nature.com/doifinder/10.1038/nature11451.

Lovett-Barron, M. et al., 2012. Regulation of neuronal input transformations by tunable dendritic inhibition. Nature Neuroscience, 15(3), pp.423–430. Available at: http://dx.doi.org/10.1038/nn.3024.

Ma, W. -p. et al., 2010. Visual Representations by Cortical Somatostatin Inhibitory Neurons--Selective But with Weak and Delayed Responses. Journal of Neuroscience, 30(43), pp.14371–14379. Available at: http://www.jneurosci.org/cgi/doi/10.1523/JNEUROSCI.3248-10.2010.

Magee, J.C., 2016. Dendritic voltage-gated ion channels. In Dendrites. Oxford University Press, pp. 259–284. Available at: http://www.oxfordscholarship.com/view/10.1093/acprof:oso/9780198745273.001.0001/acprof-9780198745273-chapter-9 [Accessed May 7, 2017].

Major, G. et al., 2008. Spatiotemporally Graded NMDA Spike/Plateau Potentials in Basal Dendrites of Neocortical Pyramidal Neurons. Journal of Neurophysiology, 99(5), pp.2584–2601. Available at: http://jn.physiology.org/cgi/doi/10.1152/jn.00011.2008.

Major, G., Larkum, M.E. & Schiller, J., 2013. Active Properties of Neocortical Pyramidal Neuron Dendrites. Annual Review of Neuroscience, 36(1), pp.1–24. Available at: http://www.annualreviews.org/doi/abs/10.1146/annurev-neuro-062111-150343.

Markram, H. et al., 2004. Interneurons of the neocortical inhibitory system. Nature Reviews Neuroscience, 5(10), pp.793–807. Available at: http://www.nature.com/doifinder/10.1038/nrn1519.

Mel, B.W., 1992. NMDA-Based Pattern Discrimination in a Modeled Cortical Neuron. Neural Computation, 4(4), pp.502–517. Available at: http://www.mitpressjournals.org/doi/10.1162/neco.1992.4.4.502.

Mel, B.W., 1993. Synaptic integration in an excitable dendritic tree. Journal of neurophysiology, 70(3), pp.1086–101. Available at: http://linkinghub.elsevier.com/retrieve/pii/S0896627313002675.

Milojkovic, B.A. et al., 2004. Burst generation in rat pyramidal neurones by regenerative potentials elicited in a restricted part of the basilar dendritic tree. The Journal of Physiology, 558(1), pp.193–211. Available at: http://www.ncbi.nlm.nih.gov/pubmed/15155788.

Mizuseki, K. et al., 2009. Theta Oscillations Provide Temporal Windows for??Local Circuit Computation in the Entorhinal-Hippocampal Loop. Neuron, 64(2), pp.267–280.

Moradi, K. et al., 2013. A fast model of voltage-dependent NMDA receptors. Journal of Computational Neuroscience, 34(3), pp.521–531. Available at: http://link.springer.com/10.1007/s10827-012-0434-4.

Müllner, F.E., Wierenga, C.J. & Bonhoeffer, T., 2015. Precision of Inhibition: Dendritic Inhibition by Individual GABAergic Synapses on Hippocampal Pyramidal Cells Is Confined in Space and Time. Neuron, 87(3), pp.576–589. Available at: http://linkinghub.elsevier.com/retrieve/pii/S089662731500625X.

Nevian, T. et al., 2007. Properties of basal dendrites of layer 5 pyramidal neurons: a direct patch-clamp recording study. Nature Neuroscience, 10(2), pp.206–214. Available at: http://www.nature.com/doifinder/10.1038/nn1826.

Palmer, L.M. et al., 2014. NMDA spikes enhance action potential generation during sensory input. Nature Neuroscience, 17(3), pp.383–390. Available at: http://www.ncbi.nlm.nih.gov/pubmed/24487231.

Pérez-Garci, E., Larkum, M.E. & Nevian, T., 2013. Inhibition of dendritic Ca 2+ spikes by GABA B receptors in cortical pyramidal neurons is mediated by a direct G i/o -βY~ subunit interaction with Ca v 1 channels. The Journal of Physiology, 591(7), pp.1599–1612. Available at: http://doi.wiley.com/10.1113/jphysiol.2012.245464.

Poirazi, P., Brannon, T. & Mel, B.W., 2003a. Arithmetic of Subthreshold Synaptic Summation in a Model CA1 Pyramidal Cell. Neuron, 37(6), pp.977–987. Available at: http://linkinghub.elsevier.com/retrieve/pii/S089662730300148X.

Poirazi, P., Brannon, T. & Mel, B.W., 2003b. Pyramidal Neuron as Two-Layer Neural Network. Neuron, 37(6), pp.989–999. Available at: http://linkinghub.elsevier.com/retrieve/pii/S0896627303001491.

Poirazi, P. & Mel, B.W., 2001. Impact of Active Dendrites and Structural Plasticity on the Memory Capacity of Neural Tissue. Neuron, 29(3), pp.779–796. Available at: http://linkinghub.elsevier.com/retrieve/pii/S0896627301002525.

Poleg-Polsky, A., 2015. Effects of Neural Morphology and Input Distribution on Synaptic Processing by Global and Focal NMDA-Spikes K. S. Brown, ed. PLOS ONE, 10(10), p.e0140254. Available at: http://dx.plos.org/10.1371/journal.pone.0140254.

Polsky, A., Mel, B. & Schiller, J., 2009. Encoding and Decoding Bursts by NMDA Spikes in Basal Dendrites of Layer 5 Pyramidal Neurons. Journal of Neuroscience, 29(38), pp.11891–11903. Available at: http://www.jneurosci.org/content/29/38/11891.long.

Pouille, F., 2001. Enforcement of Temporal Fidelity in Pyramidal Cells by Somatic Feed-Forward Inhibition. Science, 293(5532), pp.1159–1163. Available at: http://www.sciencemag.org/cgi/doi/10.1126/science.1060342.

Pouille, F. et al., 2009. Input normalization by global feedforward inhibition expands cortical dynamic range. Nature Neuroscience, 12(12), pp.1577–1585. Available at: http://www.ncbi.nlm.nih.gov/pubmed/19881502.

Rall, W. et al., 1966. Dendrodendritic synaptic pathway for inhibition in the olfactory bulb. Experimental Neurology, 14(1), pp.44–56. Available at: http://linkinghub.elsevier.com/retrieve/pii/0014488666900239.

Rall, W. & Shepherd, G.M., 1968. Theoretical Reconstruction Dendrodendritic of Field Potentials and in Olfactory Bulb Synaptic Interactions. Journal of Neurophysiology, 6(31), pp.884–915.

Rapp, M., Yarom, Y. & Segev, I., 1992. The Impact of Parallel Fiber Background Activity on the Cable Properties of Cerebellar Purkinje Cells. Neural Computation, 4(4), pp.518–533.

Rhodes, P., 2006. The Properties and Implications of NMDA Spikes in Neocortical Pyramidal Cells. Journal of Neuroscience, 26(25), pp.6704–6715. Available at: http://www.jneurosci.org/cgi/doi/10.1523/JNEUROSCI.3791-05.2006.

Royer, S. et al., 2012. Control of timing, rate and bursts of hippocampal place cells by dendritic and somatic inhibition. Nature Neuroscience, 15(5), pp.769–775. Available at: http://www.nature.com/doifinder/10.1038/nn.3077.

Salin, P. & Prince, D., 1996. Spontaneous GABA-A receptor mediated inhibitory currents in adult rat somatosensory cortex. J Neurophysiol, 75(4), pp.1573–1588.

Sanders, H. et al., 2013. NMDA and GABAB (KIR) Conductances: The “Perfect Couple” for Bistability. Journal of Neuroscience, 33(2), pp.424–429. Available at: http://www.jneurosci.org/cgi/doi/10.1523/JNEuROSCI.1854-12.2013.

Sandler, M., Shulman, Y. & Schiller, J., 2016. A Novel Form of Local Plasticity in Tuft Dendrites of Neocortical Somatosensory Layer 5 Pyramidal Neurons. Neuron, 90(5), pp.1028–1042. Available at: http://dx.doi.org/10.1016/j.neuron.2016.04.032.

Schiller, J. et al., 2000. NMDA spikes in basal dendrites of cortical pyramidal neurons. Nature, 404(6775), pp.285–9. Available at: papers3://publication/doi/10.1038/35005094%5Cnhttp://www.nature.com/doifinder/10.1038/35005094%5Cnhttp://www.ncbi.nlm.nih.gov/pubmed/10749211.

Schiller, J. & Schiller, Y., 2001. NMDA receptor-mediated dendritic spikes and coincident signal amplification. Current Opinion in Neurobiology, 11(3), pp.343–348. Available at: http://linkinghub.elsevier.com/retrieve/pii/S0959438800002178.

Segev, I. & Rall, W., 1988. Computational study of an excitable dendritic spine. Journal of neurophysiology, 60(2), pp.499–523. Available at: http://www.ncbi.nlm.nih.gov/pubmed/2459320.

Shouval, H.Z., Bear, M.F. & Cooper, L.N., 2002. A unified model of NMDA receptor-dependent bidirectional synaptic plasticity. Proceedings of the National Academy of Sciences, 99(16), pp.10831–10836. Available at: http://www.pnas.org/cgi/doi/10.1073/pnas.152343099.

Silberberg, G. & Markram, H., 2007. Disynaptic Inhibition between Neocortical Pyramidal Cells Mediated by Martinotti Cells. Neuron, 53(5), pp.735–746.

Smith, S.L. et al., 2013. Dendritic spikes enhance stimulus selectivity in cortical neurons in vivo. Nature, 503(7474), pp.115–120. Available at: http://www.nature.eom/nature/joumal/vaop/ncurrent/fuN/nature12600.html%5Cnhttp://www.ncbi.nlm.nih.gov/pubmed/24162850.

Spruston, N., Jonas, P. & Sakmann, B., 1995. Dendritic glutamate receptor channels in rat hippocampal CA3 and CA1 pyramidal neurons. The Journal of physiology, 482 (Pt 2, pp.325–52. Available at: http://www.ncbi.nlm.nih.goV/pubmed/7536248%0Ahttp://www.pubmedcentral.nih.gov/articlerender.fcgi?artid=PMC1157732.

Stokes, C.C.a, Teeter, C.M. & Isaacson, J.S., 2014. Single dendrite-targeting interneurons generate branch-specific inhibition. Frontiers in Neural Circuits, 8(November), p.139. Available at: http://journal.frontiersin.org/Journal/10.3389/fncir.2014.00139/abstract.

Takahashi, N. et al., 2016. Active cortical dendrites modulate perception. Science, 354(6319), pp.1587–1590. Available at: http://www.sciencemag.org/lookup/doi/10.1126/science.aah6066.

Vargas-Caballero, M., 2004. Fast and Slow Voltage-Dependent Dynamics of Magnesium Block in the NMDA Receptor: The Asymmetric Trapping Block Model. Journal of Neuroscience, 24(27), pp.6171–6180. Available at: http://www.jneurosci.org/cgi/doi/10.1523/JNEUR0SCI.1380-04.2004.

Villa, K.L. et al., 2016. Inhibitory Synapses Are Repeatedly Assembled and Removed at Persistent Sites In Vivo. Neuron, 89(4), pp.756–769. Available at: http://dx.doi.org/10.1016/j.neuron.2016.01.010.

Wehr, M. & Zador, A.M., 2003. Balanced inhibition underlies tuning and sharpens spike timing in auditory cortex. Nature, 426(6965), pp.442–446. Available at: http://www.nature.com/doifinder/10.1038/nature02116.

Wilmes, K.A., Sprekeler, H. & Schreiber, S., 2016. Inhibition as a Binary Switch for Excitatory Plasticity in Pyramidal Neurons C. C. Hilgetag, ed. PLOS Computational Biology, 12(3), p.e1004768. Available at: http://dx.plos.org/10.1371/journal.pcbi.1004768.

Wilson, D.E. et al., 2016. Orientation selectivity and the functional clustering of synaptic inputs in primary visual cortex. Nature Neuroscience, 19(8), pp.1003–1009. Available at: http://www.nature.com/doifinder/10.1038/nn.4323%5Cnhttp://www.ncbi.nlm.nih.gov/pubmed/27294510.

